# TCR-TRANSLATE: Conditional Generation of Real Antigen Specific T-cell Receptor Sequences

**DOI:** 10.1101/2024.11.11.623124

**Authors:** Dhuvarakesh Karthikeyan, Colin Raffel, Benjamin Vincent, Alexander Rubinsteyn

## Abstract

The paradoxical nature of T-cell receptor (TCR) specificity, which requires both precise recognition and adequate coverage of antigenic peptide-MHCs (pMHCs), poses a fundamental challenge in immunology. Efforts at modeling this complex many-to-many mapping have been greatly impeded by a severe lack of experimental data. To address this, we present TCR-TRANSLATE, a novel framework that adapts low-resource machine translation techniques to the TCR:pMHC specificity domain. Here, we explore sequence-to-sequence (seq2seq) modeling with various training strategies, including semi-synthetic data augmentation and multi-task objectives to generate antigen specific TCR sequences for a given target of interest. We benchmark twelve model variants derived from the BART and T5 model architectures on a target-rich validation set of well-studied pMHCs, finding an optimal model, TCRT5, that generated validated antigen-specific CDR3*β* sequences for previously unseen antigens. While current limitations include a narrow validation set and a focus on the CDR3*β* loop, our approach demonstrates the potential of seq2seq models in rapidly generating antigen-specific TCR repertoires, offering a promising avenue for increasing throughput in precision immunotherapies. Our findings highlight both the capabilities and limitations of sequence-based conditional TCR design, emphasizing the need for experimental validation to bridge the gaps between predictions, metrics, and functional capacity.

## 1 Introduction

Cytotoxic (CD8+) T cells are a key component of the adaptive immune system, responsible for clearing viral infections, suppressing tumor progression, and inducing a number of autoimmune conditions. They scan our bodies’ cells in the periphery, using a highly specific class of pattern recognition receptors called T-cell Receptors (TCRs) to bind complementary peptide fragments presented at the cell surface by major histo-compatibility complexes (MHCs). In this way, T-cells perform the critical function of self-nonself discrimination by accurately identifying anomalous peptide-MHCs (pMHCs), with single-amino acid precision [1]. This level of specificity makes T-cell based therapeutics an attractive treatment modality, especially in cases where a population of offending cells must be targeted without depleting the broader subtype altogether. As a therapeutic paradigm, T cell based therapies including chimeric antigen receptor T cell (CAR-T), engineered TCRs (TCR-T), and recently TCR bispecifics have led to numerous advances in the treatment of chronic infections such as CMV [2, 3] and HIV [4], autoimmune diseases [5, 6], and even solid tumors [7–10]. A critical bottleneck in the development pipeline of these cellular therapies is the identification of a TCR that is specific to foreign antigens and simultaneously tolerant to self. Current approaches use TCR discovery platforms that rely on stimulation assays and cell-sorting to identify antigen-specific clones, with high resource cost and low reported hit rates [11, 12]. Thus, *in-silico* methods to decipher the mapping between TCRs and pMHCs have the potential to transform the field of precision medicine by more readily operationalizing a 400 million year old [13] mechanism of functionally deleting cells at the sub-protein resolution.

### 1.1 Background and Related Methods

T cell effector function is carried out in a decentralized manner by a repertoire of distinct clonal subpopulations, each defined by its own stochastically generated TCR, recognizing and responding to a set of cognate pMHCs. These TCRs must therefore adequately cover the space of pathogenic pMHCs, while maintaining enough specificity to avoid self. Remarkably, this constrained optimization problem has given rise to TCR:pMHC interactions that are highly specific, but also paradoxically degenerate. Experimentally, TCRs have been shown to recognize up to 10^6^ unique peptides [14], and vice versa [15, 16]. Efforts at modeling this many-to-many mapping have been blocked by an extremely sparsely sampled TCR:pMHC cross-reactivity landscape, with most TCRs of validated specificity being identified in the context of only a few, well-studied pMHCs [17]. To date, on the order of 1% of the theoretical epitope-specific repertoire for these pMHCs has been observed.

Current approaches in modeling antigen specificity of TCRs have predominantly relied on framing TCR:pMHC cross reactivity as a binary classification task [18–33], with limited utility in TCR design [34]. Prior work exploring generative models of TCRs leveraged auto-encoders to generate realistic de novo TCRs that recapitulate repertoire level statistics in the aggregate [35, 36]. However, their use in generating epitope-specific TCR sequences is limited. More recently, [37] show the potential for reinforcement learning for TCR re-design and [38] demonstrate the potential for a 1-D CNN encoder and LSTM decoder to generate T-cell receptor sequences against a known antigen. In the structure space, ESM-1F and ProteinMPNN showed promise in recapitulating native sequences of interface residues of both CDR3*α* and CDR3*β* [39]. While we introduced an auto-regressive transformer-based architecture for this problem in [40] and related research has since emerged [41, 42], a deep understanding of the utility and limitations of conditional sequence generation methods for designing real TCRs remains elusive.

### 1.2 Previous Work

In our initial proof-of-concept [40], we explored the explicit framing of the TCR reactivity problem as a sequence-to sequence (seq2seq) task (Figure 1a). In doing so, we demonstrated the viability of our setup and models TCRBART and TCRT5, two encoder:decoder transformer models adapted from BART [43] and T5 [44] (Figure 1b), to sample antigen-specific TCR sequences conditioned on target epitope information. We introduced evaluation metrics that captured the accuracy and diversity of an epitope-specific cognate repertoire, accounting for the many-to-many mapping and data sparsity inherent to this problem. Additionally, we established the superiority of the self-attention transformer compared to parameter-matched recurrent and convolutional networks and explored the impact of decoding algorithm choice on performance. Crucially, we demonstrated the use of a target-rich validation set to better characterize true model performance and behavior by maximizing the likelihood of overlap between the model generations and the ground truth. In this work, we explore an array of training strategies, multi-task learning objectives, and data augmentation methods with the express goal of maximizing model performance, acknowledging the limitations imposed by the availability of experimental data.

**Figure 1:**
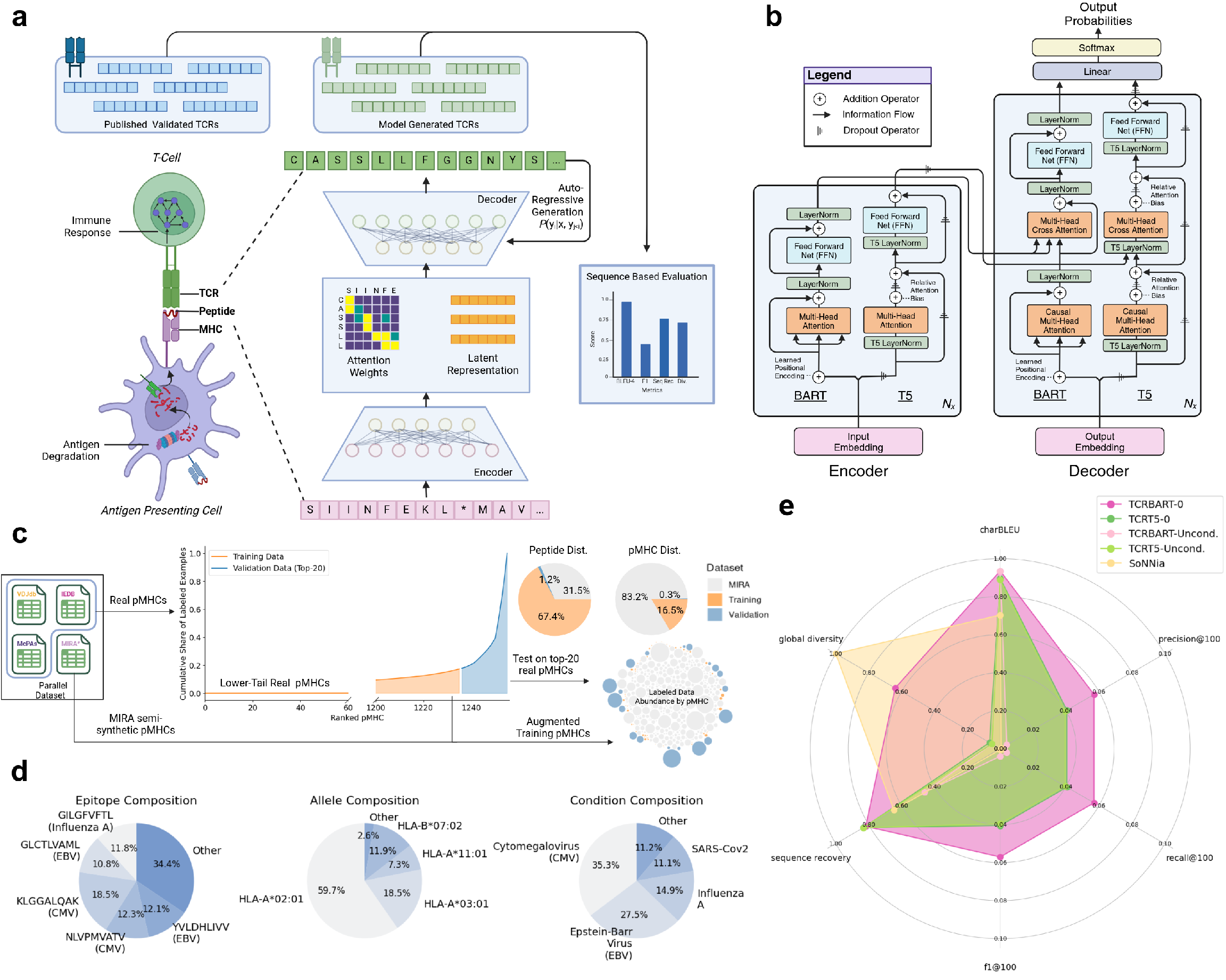
Overview of TCR-TRANSLATE. (a) Casting antigen-specific TCR design as a sequence-to- sequence (seq2seq) task. We make use of an encoder:decoder abstraction to process peptide:MHC sequence information and auto-regressively sample target-conditioned CDR3*β* sequences. (b) Specific architecture of TCRBART and TCRT5. Transformer architecture juxtaposing BART and T5 encoder and decoder layers highlighting key operations to the residual stream, inspired by (Vaswani et al., 2017). (c) Dataset creation. Given severe data-sparsity, the top-20 pMHCs from IEDB, VDJdb, and McPAS in terms of known TCRs was witheld as validation, while the remainder was used for training with semi-synthetic pMHCs from MIRA. (d) Composition of validation set. Breakdown of epitopes, alleles, and disease contexts of the top-20 real pMHCs. (e) Conditional generation outperforms unconditional generation methods. Radar plot showing the performance of TCRBART and TCRT5 models trained without pre-training (TCRBART-0, TCRT5-0) evaluated against their unconditional generations (TCRBART-Unconditional, TCRT5-Unconditional) as well as the the averaged metrics over 1000 simulations of the statistical soNNia generative model.

### 1.3 Overview

In order to directly address the issues of limited parallel data (source-target pairs) on both the training and evaluation side, we explored analogous approaches from low-resourced machine translation centered around limited and biased labeled data [45]. In this setting, numerous techniques such as back-translation [46], self-training [47], transfer learning [48], and semi-supervised approaches that leverage monolingual data have been proposed with varying degrees of success. However, these approaches can exacerbate overfitting to specific domain contexts or sequence distributions, as methods of increasing accuracy via self-consistency experience a diversity trade-off [49]. A particularly effective approach leverages the reflexive nature of sequence co-dependencies between source-target pairs by jointly learning the bidirectional mapping between source and target languages [50, 51]. These bidirectionally trained models have shown significant improvements in translation quality by sharing representations and aligning latent spaces across multiple languages [52]. However, to the best of our knowledge, these approaches have not been applied to the functional protein design domain.

Here, we use multi-task training to jointly learn both directions of TCR:pMHC specificity and evaluate its effect on the conditional generation of TCR sequences. To do so, we constructed a validation dataset comprising the top-20 pMHCs with the most experimentally validated TCRs, forfeiting their inclusion in the training data to maximize exact sequence matches during evaluation. We systematically trained twelve sequence-to=sequence model variants of TCRBART and TCRT5 using these low-resourced machine translation techniques and characterized model performance under optimal conditions. In doing so, we found an appreciable increase in native sequence recovery to known CDR3*β* binders as well as an increase in number of nonzero F1 scores across pMHCs, albeit with significantly fewer unique sequences. Interestingly, this drop in diversity was driven by a preferential sampling of highly polyspecific TCRs, shown to have broad specificity against multiple unrelated pMHCs. We observed that model performance was mostly conserved across pMHCs, unsurprisingly tied directly to the number of available reference TCRs. The observed variability in performance across different pMHC epitopes highlights the critical role of training data composition and its relationship to the test set. To demonstrate real world utility, we evaluate our flagship TCRT5 model against test antigens from last years IMMREP2023 TCR specificity competition that were unseen during training and model selection and demonstrate non-random performance on pMHCs with even a limited set of reference TCRs. Our results demonstrate the potential and pitfalls of a sequence-to-sequence modeling paradigm for the sampling of real antigen-specific TCR sequences.

## 2 Methodology

In this section, we provide an explanation of our methodology, highlighting the motivation for our choices in design, experiment, and architecture in the context of desired outcomes. Our primary goal in this study is to assess whether a seq2seq model trained on the extremely sparse paired TCR:pMHC data can generate valid TCRs for unseen antigens. Inspired by the numerous parallels, we approach TCR design as a severely low-resourced machine translation problem, leaning into the field’s terminology throughout this work. Beyond the directed target sequence “translation” of TCR sequences given a source pMHC sequence, we borrow terms such as “monolingual data” (unpaired TCR and pMHC sequences) and “parallel data” (paired TCR-pMHC examples), establishing a common vocabulary with existing work. This standardized framework streamlined our investigation of various technical aspects, from training dynamics to architectural decisions, while directly addressing the fundamental challenge of learning from extremely limited training data.

In our quest to find an optimal set of hyperparameters and model weights, we depart from the traditional structure of machine learning papers. Rather than iteratively refining a model and measuring performance gains on a well characterized benchmark, we conduct an unstructured all-vs-all comparison of model architectures and training approaches to identify configurations capable of generating valid TCRs for unseen antigens. This approach both increases our chances of finding a robust model in the space of architectures and parameters, helping identify more general model-agnostic takeaways. Through this battle-royale, we find a serendipitous discovery that revealed a potential off-ramp on the path to generalizable TCR design, influencing model selection. After choosing our final model, we validate our specific training and data choices through ablation studies to assess the contribution of these elements before characterizing our selected model’s performance through both quantitative metrics and qualitative analysis. Finally, we evaluate the model in a low-resource setting, testing its ability to generate antigen-specific TCRs for sparsely represented epitopes—a key barrier to practical application of existing TCR:pMHC specificity models.

### 2.1 Setup

Following our prior work [40], we adopted the same sequence-to-sequence (seq2seq) framework, relaxing the direction of pMHC → TCR source-target pairs to train on both directions, but evaluate on the former. To represent the TCR:pMHC trimeric complex, comprised of three sub-interactions (TCR-peptide, TCR-MHC, peptide-MHC) as a source-target sequence pair, we made a few simplifying assumptions that allowed for a more straightforward problem formulation. First, we assume a stable pMHC complex, reducing the problem to a dimeric interaction between TCR and pMHC. Second, we focus on the amino acid residues at the binding interface. For the TCR, we use the CDR3*β* loop, a contiguous span of 8-20 amino acids that typically make the most contact with the peptide [53]. Similarly, for the pMHC, we use the whole peptide and the MHC pseudo-sequence, defined in [54] as a reduced, noncontiguous, string containing the polymorphic amino acids within 4.0 Å of the peptide. We opt for a single character amino-acid level tokenization, primarily for its interpretibility [55]. In addition to the 20 canonical amino acids, we use standard special tokens including the start/end of sequence, masking, padding, and a separator token to delineate the boundary between the concatenated peptide and pseudosequence. For TCRT5, we additionally employ the use of sequence type tokens, retained from T5’s use of task prefixes [44], to designate translation direction:

*TCRBART:*

[SOS]EPITOPE[SEP]PSEUDOSEQUENCE[EOS] ↔ [SOS]CDR3BSEQ[EOS]

*TCRT5:*

[PMHC]EPITOPE[SEP]PSEUDOSEQUENCE[EOS] ↔ [TCR]CDR3BSEQ[EOS]

### 2.2 Dataset

#### 2.2.1 Parallel Corpus

As with all sequence-to-sequence tasks, we first assembled a parallel corpus between valid source and target sequences. Our parallel corpus comprised experimentally validated immunogenic TCR:pMHC pairs taken from publicly available databases (McPAS [56], VDJdb [57], and IEDB [58]). Additionally, we used a large sample of partially-labeled data derived from the MIRA [59] dataset, which contained CDR3*β* and peptide sequences, but had MHC information at the haplotype resolution instead of the actual presenting MHC allele. Therefore, the presenting MHC allele was inferred from the individual’s haplotype using MHCFlurry2.0’s [60] ranked presentation score. Of importance, these semi-synthetic examples were not used in evaluation. More on the dataset standardization procedures can be found in the Supplementary Methods (B.1).

#### 2.2.2 Training/Validation Split

To accurately assess the capacity of the models to sample antigen-specific sequences on unseen epitopes, we held out a validation set of the top-20 most target-rich pMHCs, which collectively account for over 80% of the labeled TCR sequences. We trained on the remaining data, further removing the occurrences of the held-out epitopes bound alternate MHCs to ensure a clean validation split (Figure 1c). We retained training sequences with a low edit distance to the validation pMHCs to better understand their influence on performance. The degree to which these sequences exhibit training set similarity is reflected in (Table S2). The parallel corpus was subsequently deduplicated to remove near duplicates (peptides with the same allele and a >= 6-mer overlap) which we found to marginally help overall performance, in accordance with [61]. This resulted in a final dataset split of ≈330k training sequence pairs (N=6989 pMHCs) and 68k validation sequence pairs (N=20 pMHCs). A key limitation of this dataset is its highly skewed bias towards mainly viral epitopes and a very narrow HLA distribution towards well studied alleles (Figure 1d).

### 2.2.3 Unlabeled ‘Monolingual’ Data

In addition to the experimentally validated paired data, we sought to make use of ‘monolingual’ TCR and pMHC sequences identified from single-cell V(D)J-sequencing and MHC-binding assay experiments, respectively. We hypothesized that pre-training the encoder:decoder model using a self-supervised denoising autoencoding objective could help boost the translation performance of the model by learning better representations for source and target sequences. This mode of transfer learning has been applied in the machine translation before in [62], and crucially has been shown to improve performance in the low-resource setting [63]. For the unlabeled pMHC sequences, we used the positive MHC ligand binding assay data from IEDB (N≈740K) [58]. For the TCR sequences, we used around (N≈14M) sequences from TCRdb [64] of which around 7M CDR3*β* sequences were unique. For this dataset we chose to retain duplicate CDR3*β* sequences as the TCRdb was amassed over multiple studies and populations, so we felt that the inclusion of duplicate CDR3*β*s was reflective of convergent evolution in the true unconditional TCR distribution.

### 2.3 Model Training

For our experiments, we considered six model training schemes comprising three levels of learning tasks, across pre-training status. This was done for both TCRBART as well as TCRT5 for a total of twelve model variants. All models are evaluated on CDR3*β* sequence generation (Figure 2a). The baseline models (TCRBART-0 and TCRT5-0) were trained on the pMHC → TCR direction with no pre-training. A second set of bidirectional models (TCRBART-0 (B) and TCRT5-0 (B)) were trained on conditional sequence reconstruction in both directions, and finally a set of multi-task models were trained on both directions as well a masked language modeling loss term for both TCR and pMHC sequences (TCRBART-0 (M) and TCRT5-0 (M)). Similarly six models were pre-trained and finetuned using the same learning tasks to add TCRBART-FT, TCRBART-FT (B), TCRBART-FT (M), TCRT5-FT, TCRT5-FT (B), and TCRT5-FT (M). Our model naming scheme is summarized in table form below:

**Figure 2:**
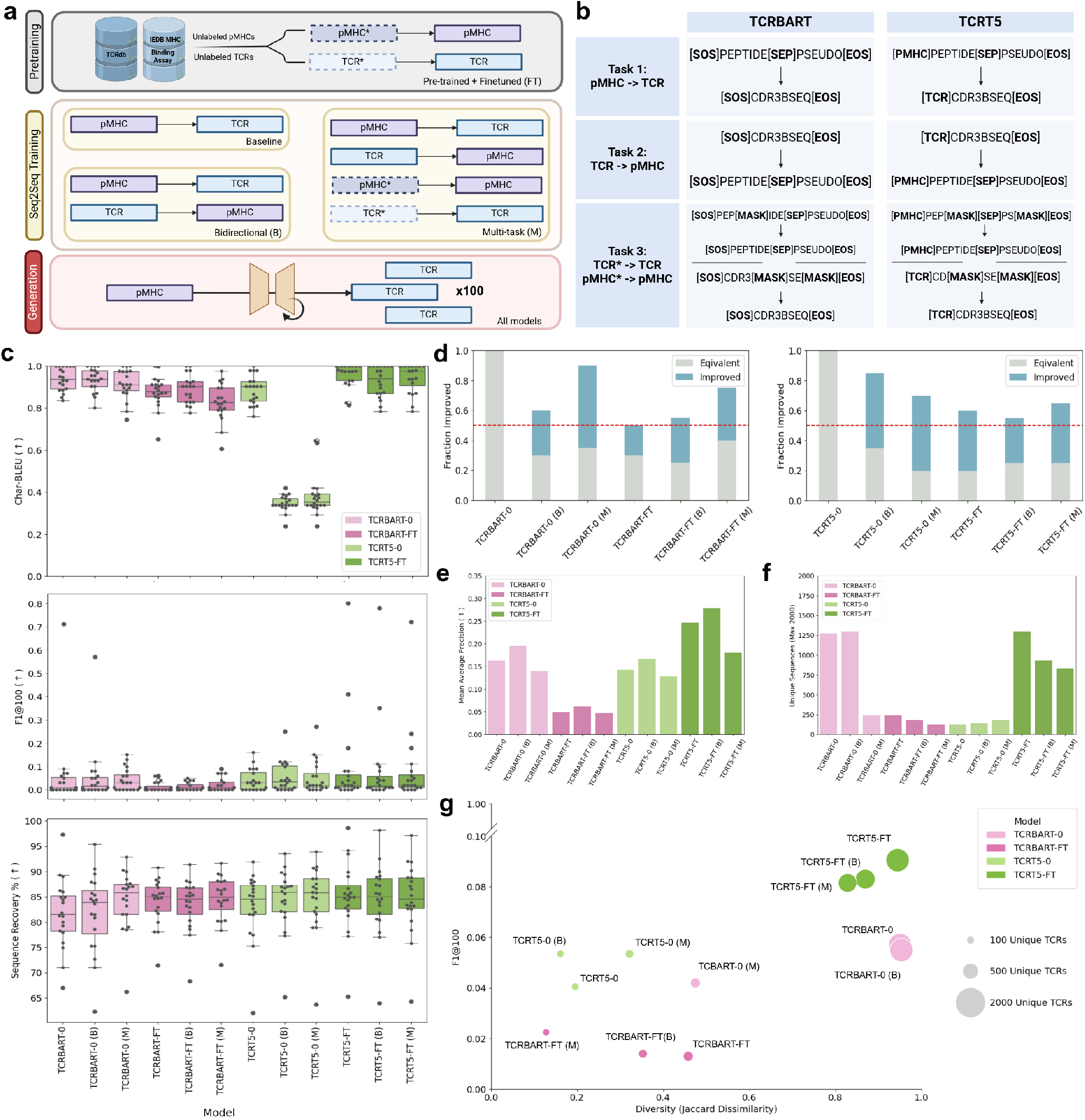
Multi-task training increases accuracy and decreases diversity metrics. (a) Diagram outlining pre-training, seq2seq training/finetuning, and their common generation scheme (inference). (b) Sequence I/O representation of TCRBART and TCRT5 broken down by task. (c) TCR-TRANSLATE accuracy metrics. Swarm plots showing the median, quartile, and individual contributions of each of the validation pMHCs for CharBLEU, F1@100, and native sequence recovery. (d) Fraction of pMHC F1@100 scores that remain equivalent to or greater than the baseline models (TCRBART-0, TCRT5-0). Red line marks the 50% point, indicating no gain in performance compared to baseline. (e) Model calibration as measured by mean average precision (mAP) across pMHCs calculated using sequence likelihood based rank per model. (f) Barplot of global diversity calculated as the total number unique sequences across pMHCs (20 × 100=2000 max). (g) Scatterplot summarizing model performance on accuracy and diversity metrics. Accuracy is taken as the mean F1@100 score and diversity is shown both in terms of the total number of unique sequences generated (size of each data point) as well as the mean pairwise Jaccard dissimilarity scores across pMHCs (x-axis).

|  | Pre-training Status |  | Training Method |  |  |
| --- | --- | --- | --- | --- | --- |
|  | No | Yes | Baseline | Bidirectional | Multi-task |
| TCRBART | -0 | -FT | — | (B) | (M) |
| TCRT5 | -0 | -FT | — | (B) | (M) |

#### 2.3.1 Pre-Training

In addition to their architectures, the performance of the original BART and T5 models can be attributed to their pre-training objectives, as shown in their respective papers. However, some objectives, like document rotation in BART, are less applicable for biological sequences. Therefore, we focus on masked denoising objectives commonly used in protein language models. TCRBART was pre-trained using masked amino acid modeling (BERT-style [65]), while TCRT5 utilized masked span reconstruction, learning to fill in randomly dropped spans with lengths between 1 and 3. Of importance, neither model was trained on complete sequence reconstruction to reduce the possibility of memorization during pre-training. Both models were trained on unlabeled CDR3*β* and peptidepseudosequences, simultaneously pre-training the encoder and decoder, inspired by the MASS/XLM approach [66, 67]. Unlike MASS/XLM, we omitted learned language embeddings, allowing the model to learn from the size differences between CDR3*β* and pMHC sequences. To address the imbalance in sequence types, we upsampled sequences for a 70/30 TCR to pMHC split.

#### 2.3.2 Direct Training/Fine-tuning

For the parallel data, we used the same three training regimes (baseline, bidirectional, multi-task) for direct training from random initialization as well as finetuning. This was done by extending the standard categorical cross entropy loss function (Equation 1), favored in seq2seq tasks for its desired effect of maximizing the conditional likelihoods over target sequences [68, 69]. For the baseline training, we used the canonical form of the cross entropy loss, as shown below:

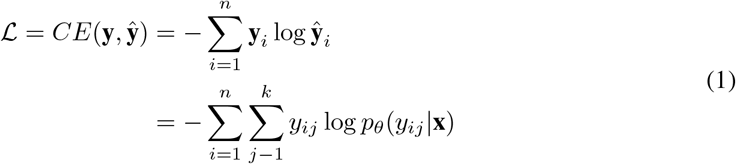

The bidirectional and multi-task models were trained using mutli-term objectives, forming a linear combination of individual loss terms corresponding to the cross entropy loss of each task/direction.

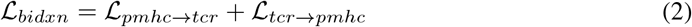

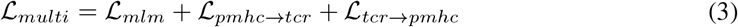

In order to mitigate effects of model forgetting with stacking single-task training epochs, we shuffled the tasks across the epoch using a simple batch processing algorithm (Algorithm 1). After the batch was sampled, it was rearranged into one of four sequence-to-sequence mapping possibilities and trained on target reconstruction with the standard cross entropy loss, which was used for back-propagation. In this way, we could ensure that the model was simultaneously learning multiple tasks during training. For the bidirectional model, this was straightforward as we could swap the input and output tensors during training to get the individual loss contributions of the ℒ_*pmhc*→*tcr*_ and ℒ_*tcr*→*pmhc*_ (Equation 2). For the multi-task model, the mapping possibilities are: 1) pMHC → TCR 2) pMHC → TCR 3) Corrupted pMHC* → pMHC 4) Corrupted TCR* → TCR, which combine to to form ℒ_*multi*_ (Equation 3). These tasks and sequence mappings as seen by TCRBART and TCRT5 are summarized in Figure 2b.

##### Algorithm 1

Multi-Task Training Step

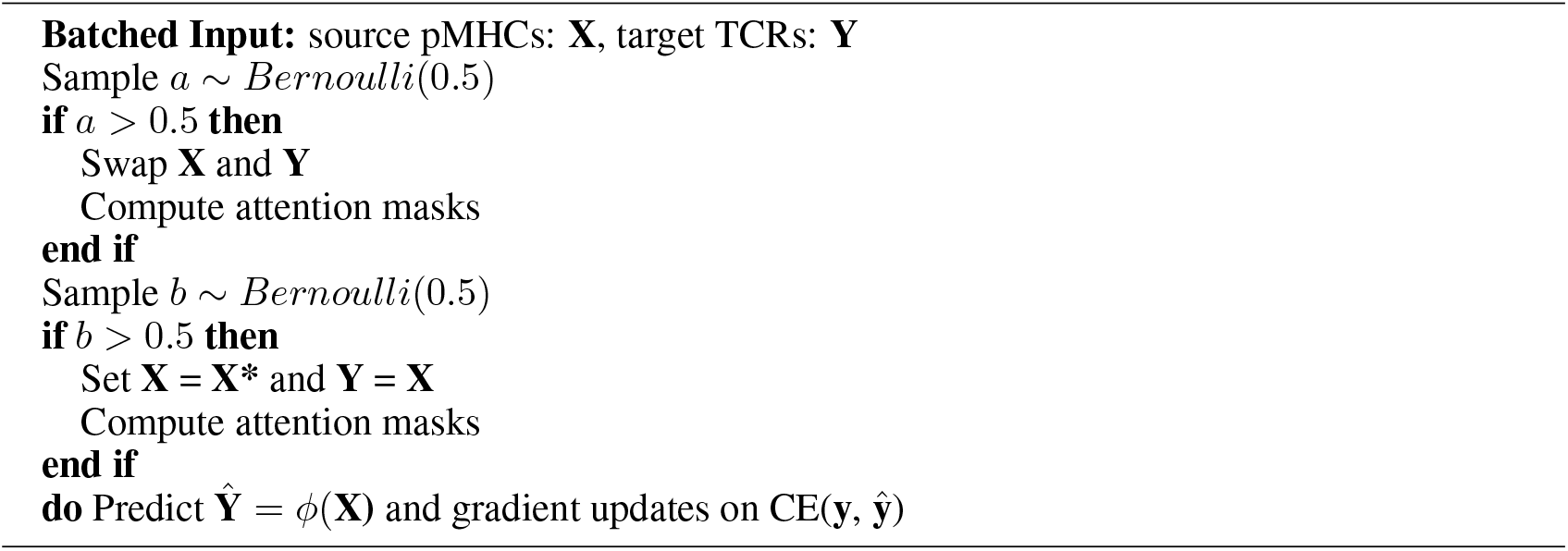

For the purposes of comparison between models originating from different training schemes, each of the models was trained for 20 epochs, from which the checkpoint with the highest average overlap to the known TCR reference set (F1 score) was chosen. We chose this approach to characterize the models’ real world potential under optimal conditions, as opposed to training for a fixed number of steps or even a fixed number of steps per task (See B.3 for Checkpoint Selection).

### 2.4 Generation Algorithm

Given its marked performance in the NLP setting, we selected beam search for all sequence generations, except for calculating the dataset-level Char-BLEU, where we used greedy decoding in line with previous work. Both decoding methods and the broader class of mode-seeking methods, aim to maximize for the highest probability conditional target sequence. A desirable property of mode seeking algorithms is that they are deterministic. Recently, however, mode-seeking algorithms have come under scrutiny for sampling only a small portion of the true target distribution, as noted in [70]. Instead of sampling tokens directly from the whole conditional distribution: *y*_*t*_ ∼ *P* (*y*_*t*_|*y*_*<t*_, *x, θ*), given source sequence *x* and model parameters *θ*, mode-seeking algorithms try to approximate the *y*^*MAP*^ = arg max_*y*∈*Y*_ log *p*(*y*|*x, θ*) which explores the conditional distribution about the mode. However, combined with the ability to assess and maintain longer high probability subsequences, the use of beam search results in a powerful method for sampling native sequence-like predictions, which we observed to be true for the TCR space in our prior work [40].

### 2.5 Evaluation

Evaluating the performance of generative models for biological sequences, such as TCR sequences, poses unique challenges. Due to the high cost of *in-vitro* validation, assessing the fidelity and usefulness of generated sequences is difficult, particularly for de novo sequences. When even the most well represented pMHCs have on the order of 10^4^ experimentally validated TCRs out of the theoretical 10^6^, traditional accuracy and recall-based metrics often low-ball performance, inadvertently testing similarity to an arbitrary sample of validated examples (Figure S1).

In natural language processing, model-based scores such as BLEURT [71] and COMET [72] have been used as a means of scoring generated outputs when faced with data constraints. While one could use existing TCR-epitope recognition predictors to evaluate the antigen-specificity of the models’ generations, their known issues with out of distribution generalization [17, 73, 74] risk introducing an opaque bias that is additionally confounded by these models’ varying training data. In the antibody space, binding affinity between the antibody and the target is known to be one of the critical determinants of functional activity [75]. As such, accurate methods of *in-silico* binding affinity prediction have been used to evaluate and even optimize de novo antibodies on their antigen-specificity [76, 77]. However, such methods of *in-silico* evaluation of TCR:epitope specificity are less meaningful, since unlike antibodies, binding affinity and structural fit alone do not predict functional response of TCRs [78–80].

To evaluate antigen-specificity, we build our framework around sampling exact CDR3*β* sequences from published experimental data on well-characterized validation epitopes not seen during training. This approach has an interpretable bias compared to black-box error profiles, at the cost of potentially underrepresenting actual performance. We calculate sequence similarity-based metrics beyond exact overlap to create a more robust evaluation framework, and characterize their concordances for future use on epitopes with fewer known cognate sequences. Broadly, our metrics can be summarized as evaluating the accuracy of the returned sequences, their diversity, or some combination of the two. They are summarized in brief below:

#### Accuracy Metrics

- **Char-BLEU**: Following BLEU-4 [81], the character-level BLEU calculates the weighted n-gram precision against the *k* = 20 closest reference sequences to abate unintended penalization of accurate predictions under a large reference set. We use the NLTK’s `‘sentence_bleu’` function to calculate a single translation’s BLEU score and the `‘corpus_bleu’` function to compute the BLEU score over an entire dataset.
- **Native Sequence Recovery**: We compute the index-matched sequence overlap with the closest known binder of the same sequence length, when available. This is the same as the length-normalized Hamming distance. The Levenshtein distance normalized to the length of closest reference was used for cases where a size-matched reference did not exist.
- **Mean Average Precision (mAP)**: Borrowed from information retrieval, mean average precision measures the average precision across the ranked model predictions. Here, we rank the generations by model log-likelihood scores and take the average of the precisions at the top-1, top-2, top-3, … top-k ranked outputs. Then we take the mean over the various pMHCs’ average precision (AP) values to get the mean average precision. This metric gauges both the accuracy of the model as well as the calibration of its sequence likelihoods.
- **Biological Likelihood**: To assess the plausibility of model outputs independent of antigen-specificity or labeled data, we compute generation probability of predictions using OLGA, a domain specific generative model that infers CDR3*β* sequence likelihood [82].

#### Diversity Metrics

- **Total Unique Sequences**: As a measure of global diversity, we compute the number of total unique generations across the top-20 validation pMHCs as a diversity metric that captures model degeneracy and input specificity.
- **Jaccard Similarity/Dissimilarity Index**: The Jaccard Index or the Jaccard similarity score is used to measure the similarity of two sets and is calculated as the size of the intersection divided by the union of the two sets. Since the Jaccard Index is inversely proportional to diversity, one minus the Jaccard Index is often to use to represent diversity between two sets.
- **Positional** Δ**Entropy**: In order to quantify the change in diversity between the models’ outputs and the reference distribution per CDR3*β* position, we report *H*(*q*_*i*_)−*H*(*p*_*i*_) over the KL divergence to get a signed change in entropy between the amino acid usage distribution of reference distribution *q* and sample distribution *p* at position *i*.

#### Both

- **Precision, Recall, and F1@K**: Also taken from information retrieval, these metrics gauge precision, recall, and F1 by sampling *K* = 100 times, without rank, and measuring the exact sequence overlap with the reference sequences. In the case of beam search decoding, since we observed beam search to return unique sequences at our choice of decoding parameters, all of these metrics were equivalent and are simply represented by the F1 score.
- **K-mer Spectrum Shift**: As used in the DNA sequence design space [83], the k-mer spectrum shift measures the Jensen-Shannon (JS) divergence between the k-mer usage frequency distributions of two sets of sequences across different values of k. Here we compare the JS divergence between the distribution of k-mers derived from a pMHC’s model generations and its reference set of sequences.

## 3 Results

### 3.1 Conditional generation outperforms unconditional generation

Before comparing our various training techniques, we first sought to calibrate our metrics by bench-marking conditional models *P* (*TCR*|*pMHC*) against unconditional generation *P* (*TCR*). We evaluated our baseline model variants TCRBART-0 and TCRT5-0 on a reduced set of metrics to determine the express advantage of conditioning on the input over unconditional generation. As our main unconditional baseline we used soNNia’s `‘Ppost’` [36], a generative model that extends V(D)J recombination to include thymic selection, sampling a TCR distribution that is closer to what is observed in the periphery. In addition, to investigate the effects of our training data on validation performance, we sampled TCRBART-0 and TCRT5-0 in an input-free manner (TCRBART-Unconditional, TCRT5-Unconditional) to generate CDR3*β*s sequences with no pMHC information. To accommodate variability in soNNia’s stochastic generations, we sampled 100 CDR3*β*s for each of the top-20 validation pMHCs and took the average over each of our metrics from 1000 simulations. For the unconditional TCRBART and TCRT5, given the deterministic nature of beam search, we used the same set of 100 unconditional sequences for all pMHCs so as to not inject bias by way of arbitrating sequences to different pMHCs. Unsurprisingly, we found that the conditional models outperformed unconditional approaches across all metrics except global diversity (Figure 1e). TCRBART-0 showed greater performance discrepancy between conditional and unconditional modes due to degenerate sampling of the same amino acids at the C-terminus, while TCRT5-0’s difference was more subtle. In fact, TCRT5-Unconditional achieved non-zero F1 scores, indicating the presence of high-likelihood training sequences in the test set.

### 3.2 Multi-task training increases accuracy metrics while decreasing diversity metrics of generated sequences

Having established unconditional generation as the performance lower bound of conditional generation, we proceeded to benchmark the performance of the twelve model variants on our accuracy and diversity metrics. We approached the results of our model comparison by first assessing the model classes (BART and T5) separately, then across their pre-training status, and finally juxtaposing the three different tasks (baseline, bidirectional, multi-task) within each model:pre-training combination.

Under our evaluation framework, no models or training strategies consistently outperformed others on all metrics across all pMHCs (Table 1). In many instances, improvements in one metric were offset by declines in another. For example, while mean and median sequence recoveries increased for bidirectional and multi-task variants, Char-BLEU scores decreased (Figure 2c). We observed a polarization in the F1 trends, where some models excelled on a small subset of examples and others showed marginal improvements across a broader set. This was observed as a decrease in mean F1 score but an increase in median compared to the baseline models. To check if adding training complexity increased performance over baseline, we analyzed how many individual pMHCs had an improved F1@100 score. We found that all of the training procedures maintained or improved F1 performance for over 50% of validation pMHCs, indicating material benefit over baseline training (Figure 2d). Looking at the calibration of these models’ predictions to see if the real binders had higher model likelihoods as compared to the those that were unconfirmed, we scored the models on mean average precision (mAP) by ranking the predictions by their output scores and computing the average precision down the list of rankings. We found that across the board, the bidirectional models outperformed the baseline models, and both outperformed the multi-task variants (Figure 2e). However, diversity metrics revealed a decline in unique sequences generated across pMHCs going from the baseline models to the bidirectional or multi-task ones. This was most evident for TCRBART-0 (M), which had strong accuracy metrics despite a drop of over 80% in unique sequences generated (Figure 2f). Taken together, we find examples of strong model performance where accuracy metrics were high with a small number of unique sequences and examples where performance is high with a large number of unique translations, indicating that these metrics are more useful when taken together.

**Table 1:** Performance Metrics on Top-20 Validation pMHCs. Mean/Median values are reported where applicable. Best in class metric is highlighted in bold.

| Model | Char-BLEU (↑) | F1@100 (↑) | %Rec. (↑) | mAP (↑) | $N_{unique}$ (↑) |
| --- | --- | --- | --- | --- | --- |
| TCRBART-0 | **93.5** | **.057**/.010 | 81.7/81.5 | 0.163 | 1276 |
| TCRBART-0 (B) | 93.0 | .055/.015 | 81.9/83.9 | **0.196** | **1302** |
| TCRBART-0 (M) | 91.7 | .042/**.030** | **84.3/85.9** | 0.140 | 240 |
| TCRBART-FT | 87.4 | .013/0.00 | 84.1/85.1 | 0.049 | **240** |
| TCRBART-FT (B) | **89.8** | .014/**.010** | 83.5/84.6 | **0.062** | 185 |
| TCRBART-FT (M) | 86.1 | **.023/.010** | **84.5/85.0** | 0.048 | 127 |
| TCRT5-0 | **89.5** | .040/.030 | 83.2/84.7 | 0.142 | 129 |
| TCRT5-0 (B) | 34.8 | **.054/.035** | 84.5/**85.9** | **0.167** | 139 |
| TCRT5-0 (M) | 37.8 | **.054**/.020 | **84.7/85.9** | 0.129 | **177** |
| TCRT5-FT | **96.4** | **.091/.020** | **84.9**/85.0 | 0.246 | **1300** |
| TCRT5-FT (B) | 93.5 | .083/.015 | 84.6/**85.1** | **0.279** | 933 |
| TCRT5-FT (M) | 94.4 | .082/**.020** | **84.7**/84.6 | 0.180 | 833 |

In order to holistically characterize all models, we visualized their performance on a biaxial plot of both accuracy (average F1 score) and diversity (mean pairwise Jaccard dissimilarity). We chose the average F1 score as opposed to median or fraction improved for the reason that designing therapeutic TCRs requires an oversampling for the subsequent pruning of self-reactive motifs and thus a model that generates many correct sequences may be more useful in this setting. Viewed through this lens we found that all variants from both TCRBART-0 and TCRT5-0 performed roughly the same from an accuracy standpoint, except that TCRBART-0 was able to generate many more unique translations, on par with TCRT5-FT. While pre-training and finetuning pushed the diversity/accuracy pareto front for the TCRT5-FT variants, we observed the complete opposite effect of degradation in both model performance and diversity for all TCRBART-FT variants (Figure 2g, S2a-b). Since both TCRBART-FT and TCRT5-0 generated less than 10% of the maximum number of unique sequences, with average Jaccard dissimilarities of less than 0.5, we find that the variants from TCRBART-0 and TCRT5-FT to be the best BART and T5 models, with the TCRT5-FT versions achieving the higher performance metrics between the two. However, given the discrepancies in fraction improved, mAP, and native sequence recoveries, the differences between the baseline, bidirectional, and multi-task models of TCRBART-0 and TCRT5-FT were less obvious. Crucially, however, remained the fact that these bidirectional and multi-task model variants that generated far fewer sequences were still observed to improve or at least not worsen the performance of more than half the validation set. When we examined the real sequences, we saw that these models were sampling empirically de-risked TCRs that were known binders to numerous pMHCs, including those in the validation set (Figure S4).

### 3.3 Multi-task models preferentially sample polyspecific CDR3*β*equences via training set statistics

While some level of TCR cross-reactivity is an essential component that shapes the TCR repertoire, the idea of a distinctly polyspecific TCR [84], describes TCRs that bind sufficiently unrelated pMHCs (Figure 3a). Though debated as something greater than a sampling artifact [85], more recent work [86] has posited a potentially mechanistic explanation for the generation of these broadly specific TCRs as well as an evolutionary justification for their existence to maintain durable immunity. Thus, given the improbable nature in which the multi-task models maintained competitive performance with a fraction of the total unique sequences as well as their generation of sequences that were found to respond to multiple validation pMHCs, we checked the translations’ polyspecificity status as described in the literature to find the determinants that caused them to be generated multiple times across the pMHCs.

**Figure 3:**
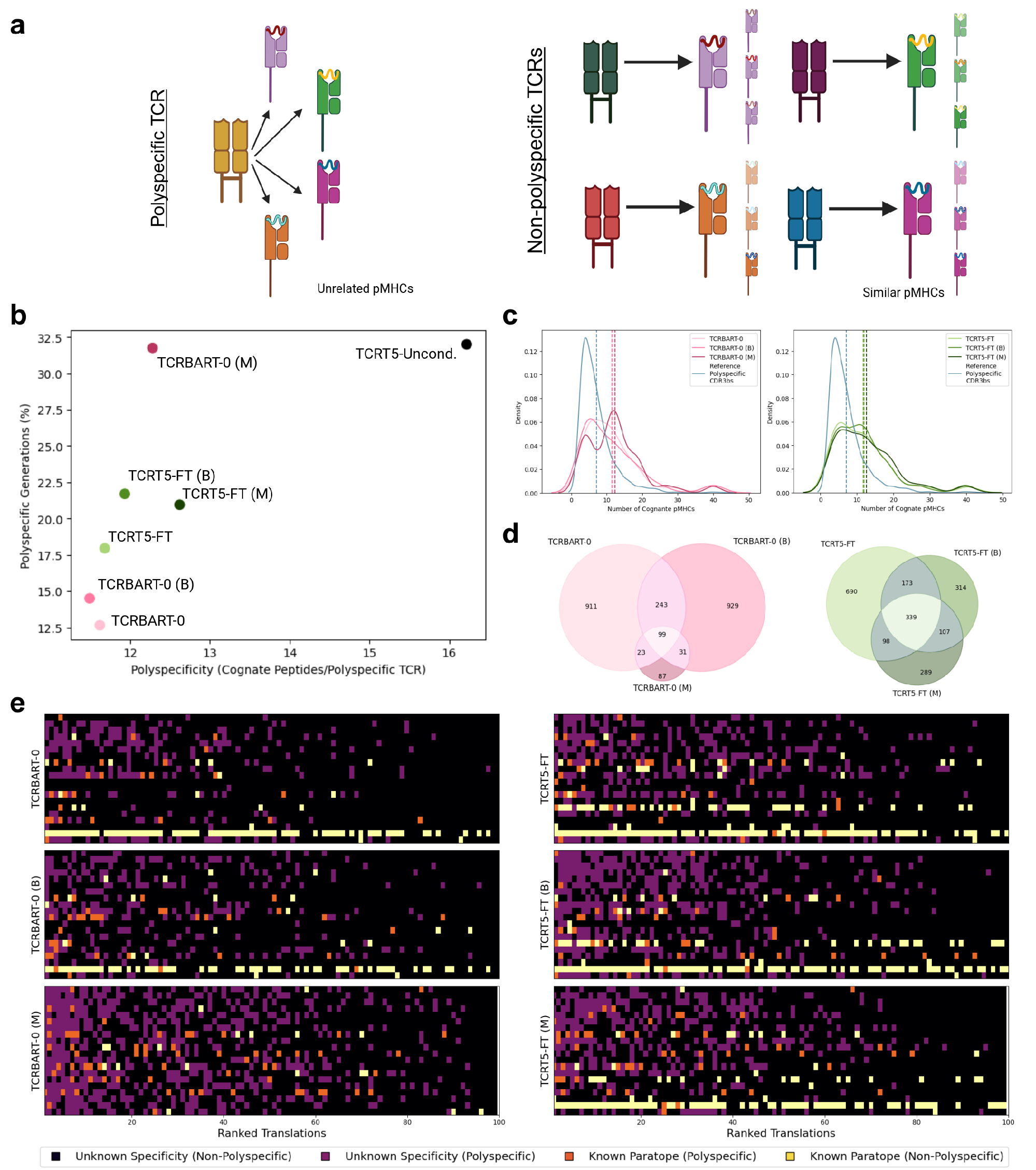
Multi-task training promotes degenerate sampling of polyspecific TCRs. (a) Diagram of showing polyspecific TCRs binding different, unrelated pMHCs juxtaposed against regular TCRs sharing a more conserved cross reactivity profile. (b) Scatterplot showing the number of polyspecific generations as a percentage as well as the mean polyspecificity (number of distinct peptides) of the polyspecific TCRs per model is shown. (c) Distribution of TCR polyspecificity across the parallel data and model generations. Density plot of cognate peptide counts for polyspecific TCRs aggregated from the combined training and validation set (reference CDR3*β*s) and the model variants per class. (d) Venn diagrams of translation overlaps for TCRBART-0 and TCRT5-FT model variants. (e) TCRBART and TCRT5 sample polyspecific and known binders with higher sequence likelihoods than those of unknown specificities. Discrete heatmaps for TCRBART-0 and TCRT5-FT variants showing the impact of increasing task complexity on the type and validity of generations.

In [86], the authors lay out the criteria for a polyspecific TCR as: (i) possessing higher probabilities of generation (ii) sampling particular V and J genes at a higher rate (iii) shared CDR3 sequences between individuals, and (iv) activation by multiple unrelated peptides. In our case, our data did not have individual level granularity, nor did it always contain V and J-gene usage. As such, we separate out a set of sequences from the combined train and test data that were found in more than one disease condition and bound more than two epitopes (n=915 CDR3*β* sequences) and checked to see if they were enriched in the multi-task translations. For this set of analyses, we analyze the models that sampled a sufficient number of unique sequences (TCRBART-0 and TCRT5-FT model families). Among these models, we found that not only did the multi-task models generate more polyspecific CDR3*β* sequences for both TCRBART-0 (*p*_*bidxn*_ = 0.048, *p*_*multi*_ *<* 0.0001) and TCRT5-FT (*p*_*bidxn*_ = 0.002, *p*_*multi*_ = 0.009), but that their mean polyspecificity as measured by their number of cognate epitopes increased as well compared to the single task models (Figure 3b). In fact, the propensity to sample polyspecific TCRs varied inversely with the number of total unique sequences generated (Pearson’s R: -.957).

However, given our dataset’s deduplication step, these polyspecific CDR3*β* sequences were also more represented than those with fewer known binders. To determine if the models were learning that these polyspecific TCRs were innately valid translations for multiple unrelated peptides, or were parroting the repeated sequences seen during training at similar frequencies, we examined the translations’ rank against their polyspecificity, number of distinct epitopes/alleles bound, and training set incidence (Figure S5a-b). We found confidently sampled sequences were more common in the training set and targeted more epitopes. Interestingly, models showed a slight trend toward sampling TCRs with more dissimilar cognate epitopes, suggesting robustness in capturing polyspecificity. Regression analysis of occurrence across validation pMHCs and training frequency showed that this proposed effect was surprisingly more attenuated in the multi-task models with Pearson’s correlation coefficients of 0.41, 0.42, and 0.3 for TCRT5-FT, TCRT5-FT (B), and TCRT5-FT (M), respectively. This trend held for TCRBART-0 with a correlation coefficients of 0.43, 0.35, and 0.2 for the baseline, bidirectional, and multi-task variants as well (Figure S5c). These results suggest that the multi-task models are sampling highly cross-reactive, polyspecific CDR3*β* sequences, somewhat independent of their training set frequency.

Since our validation set is comprised of mostly viral peptides known to be immunogenic and the targets of polyspecific TCRs [86], we assessed whether our performance on F1 could be explained solely by polyspecific generations. Our analysis revealed several of the models’ translations across pMHCs were polyspecific and that these polyspecific generations were ranked high in terms of model likelihood scores (Figure 3e). Many of these polyspecific sequences were known binders to the validation pMHCs, however, many known binders were also non-polyspecific. Given our desire for a model that generates CDR3*β* sequences to rare epitopes that likely do not have convergently evolved, high probability TCRs, we find polyspecific TCR generation a potentially misleading avenue for metric hacking, misrepresenting true usefulness. Thus, while the bi-directional and multi-task models show promise in increasing accuracy through self-consistency for the receptor:ligand design problem, we note that their utility may be limited given the current dataset distribution of asymmetrically sampled pMHCs and cognate TCRs. We therefore select the vanilla TCRT5-FT as our flagship model for its superior accuracy, diversity, as well as minimal polyspecific TCR generation, and henceforth refer to it simply as TCRT5.

### 3.4 TCRT5 Data Ablation Study

To evaluate the impact of specific training decisions, we conducted an ablation study by removing key complexities of our training and data pipelines and measuring their effects on model performance. We started with our chosen model, TCRT5, finetuned on the single-task TCR generation with semisynthetic MIRA [59] data. Next, we retrained the model without the MIRA data for an equivalent number of steps to assess its contribution. Finally, we removed pre-training altogether, training a model on the reduced dataset from random initialization.

To avoid over-representing the performance of the model trained on MIRA data on similar validation examples, we specifically removed three pMHCs that were a single edit distance from a MIRA example with a greater than 5% overlap in their cognate CDR3*β* sequences (LLLDRLNQL, TTDPSFLGRY, YLQPRTFLL) from the validation set. For all models, we used the same checkpoint heuristic, selecting the model with the highest F1 score. The results of these ablations are summarized in Table 2.

**Table 2:** Ablation Study. Mean/Median values are reported where applicable. Best in class metric is highlighted in bold. Models were evaluated on 17 validation pMHCs (max 1700 unique sequences).

| Model | +MIRA | Char-BLEU (↑) | F1@100 (↑) | %Rec. (↑) | mAP (↑) | $N_{unique}$ (↑) |
| --- | --- | --- | --- | --- | --- | --- |
| TCRT5-FT* | + | **97.1** | **.050/.020** | **83.8/84.9** | **0.207** | 1044 |
| TCRT5-FT | - | 90.1 | .027/.010 | 72.4/74.2 | 0.168 | **1666** |
| TCRT5-0 | - | 79.4 | .044/0.00 | 65.1/65.6 | 0.170 | 1060 |

We observed for the most part, each subsequent ablation resulted in a strictly worse performance with a few notable exceptions. First, the inclusion of the MIRA dataset caused a significant reduction in sequence diversity, as the TCRT5-FT (-MIRA) model, trained on a smaller but more diverse dataset, generated 1957 unique sequences out of 2000. Second, transitioning from the unconditional to the conditional model reduced Char-BLEU and sequence recovery metrics, highlighting the reasonable performance that can be achieved simply by sampling around the global mode, with the reference CDR3*β* sequences in our validation set.

### 3.5 TCRT5 generates real unseen antigen-specific CDR3*β* sequences

Having selected TCRT5 as our flagship model, exhibiting more input sensitivity and relying less on polyspecific TCRs, we proceeded to assess its usefulness in a more qualitative manner. To evaluate how well TCRT5’s translations reflected the global statistics of the reference set, we examined the distributions of CDR3*β* lengths and generation probabilities (Figure 4a). TCRT5 captured CDR3*β* lengths ranging from 10-20 with a slight decrease in diversity (mean: 14.6, sd: 1.2) compared to the reference set (mean: 14.5, sd: 2.0). However, the generations had a significantly higher log generation probability (mean: -7.04, sd: 0.85) than the reference set (mean: -9.83, sd: 2.356), indicating TCRT5 was missing the range of lower probability sequences. This contraction in TCR diversity was corroborated by various sequence embedding models, which showed that TCRT5 translations clustered in a specific region, while random and reference sequences were more dispersed (Figure S6a-c). To investigate whether this effect stemmed from beam search decoding rather than model weights, we compared generation probabilities of reference CDR3*β*s with beam search and ancestral translations. Ancestral samples spanned the reference distribution, whereas beam search sequences mostly occupied the right tail (Figure S7a). Interestingly we observed these *p*_*g*_*en* values to correlate with the model log-likelihood scores for many pMHCs (Figure S7b).

**Figure 4:**
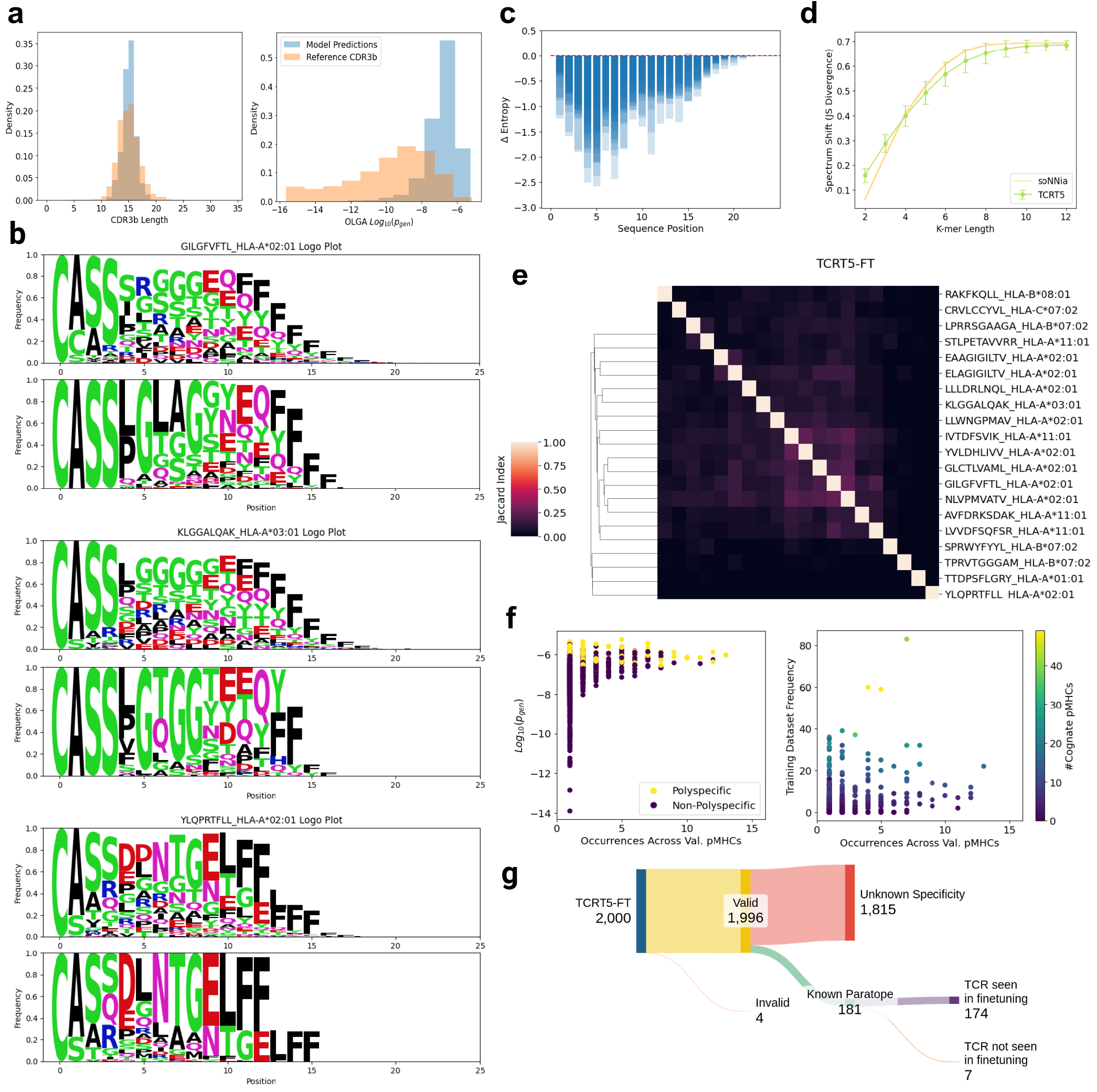
Qualitative assessment of TCRT5. (a) Repertoire level features of reference and generated CDR3*β*s. TCRT5 captures the tails of the CDR3*β* length distribution but preferentially samples sequences at the right tail of OLGA generation probabilities. (b) Sequence logo plots showing the decrease in sequence diversity position across the generated and reference CDR3*β* sequences for three canonical pMHCs [GILGFVFTL (Influenza-A), KLGGALQAK (EBV), YLQPRTFLL (SARS-CoV2)]. (c) Generated sequences experience a decrease in Shannon entropy for nearly all positions compared to reference sequences across all pMHCs. Barplots for individual pMHCs are overlaid on one another. (d) K-mer spectrum shift plot showing the Jensen Shannon divergence between generated and reference sequences. Error bars mark the mean and 1-standard deviation across validation pMHCs. (e) Heatmap of Jaccard Index scores showing the generated sequence co-occurrence across different pMHC pairs. Mean JS divergence for soNNia generations for 100 sequences sampled per pMHC across 100 simulations are shown for reference. (f) TCRT5 repeats sequences across pMHCs in line with biological probabilities and is robust to training set abundance. Scatterplot visualizing the occurrence across pMHCs with the OLGA *p*_*gen*_, polyspecificity, and training set frequency. (g) TCRT5 generates experimentally validated antigen-specific CDR3*β* sequences unseen during training. Sankey diagram showing the validity (denoted as non-zero OLGA *p*_*gen*_), known antigen-specificity status, and training set membership of generated sequences across the validation pMHCs are shown.

We then assessed how the loss of diversity affected translations for the validation pMHCs, focusing on canonical epitopes such as GILGFVFTL (Influenza A), KLGGALQAK (CMV), and YLQPRTFLL (SARS-CoV2) which have 8083, 12660, 1636 reference CDR3*β*s respectively. Sequence logo plots revealed a noticeable decrease in diversity, particularly at the sequence start due to a strong preference for sequences beginning with ‘CASS’ (Figure 4b). This loss of entropy was quantified using the positional ΔEntropy value, which showed a loss in diversity for all sequence positions across all pMHCs with the greatest loss in entropy in around position 5 (Figure 4c). Additionally, we evaluated TCRT5’s ability to match arbitrary k-mers frequencies with the reference sequences, using Jensen-Shannon divergence to assess spectra shifts. Compared to randomly generated sequences, we observed a lower divergence in soNNia generations at lower k-mer lengths, followed by an intersection, and a region where TCRT5 maintains a lower spectrum shift than the unconditional generations (Figure 4d). To account for the impact of sampling only 100 pMHCs, we also sampled 1,000 sequences and found consistent results (Figure 8).

Next, to determine TCRT5’s sensitivity to input, we computed the Jaccard index to assess overlap between translation sequences across pMHCs (Figure 4e). As expected, sequences with high similarity clustered together, such as melanoma antigens EAAGIGILTV and ELAGIGILTV. Perhaps more interestingly, we found a cluster of high overlap between dissimilar EBV epitopes, indicating that translations may be conserved by allele or disease context, though more work is required to determine to what extent this is true. To see check for correlates of sequence occurrence across validation pMHCs, we compared generation probabilities, polyspecificity, and training set frequency. Higher generation probability sequences were more frequently sampled, though no clear correlation existed between training frequency and increased sampling (Figure 4f).

To see if any of the true positive sequences were not seen during training we stratified the validity (OLGA *p*_*gen*_ *>* 0), known specificity, and training set membership of each of the 100 × 200 = 2,000 input:translation pairs and found that of the 2,000 generations, 1,996 of them had nonzero generation probabilities, 181 were known binders, and 7 pairs had TCRs that were not seen during the supervised training (Figure 4g). Notably, one of these seven (CSARDRLAQNTGELFF) was not found in the pre-training set, and had a *p*_*gen*_ = 1.68 * 10^−10^, indicating a level of robustness from simply sampling from the unconditional *P* (*TCR*) distribution). Moreover, these pairs spanned four different pMHCs: KLGGALQAK (CMV), LLWNGPMAV (YFV), YLQPRTFLL (SARS-CoV2), and YVLDHLIVV (EBV), revealing that the performance wasn’t localized to a single pMHC.

### 3.6 TCRT5 achieves non-random performance on sparsely validated epitopes

The primary goal of a TCR design model like TCRT5 is to sample TCRs against rare epitopes not seen during training, especially when few or no known TCRs exist. As highlighted in the recent IMMREP2023 TCR specificity competition, models for binary prediction often struggle to outperform random predictors in this regime [74]. We sought to evaluate TCRT5 in this context, by generating CDR3*β* sequences for “unseen” epitopes from the IMMREP test dataset (FTDALGIDEY, SALPTNADLY, TSDACMMTMY) and another epitope (TDLGQNLLY) that was absent from our training and validation data, all presented by HLA-A*01:01 (Figure 5a). When generating 100 CDR3*β* sequences, TCRT5 did not share any concordance between translation and reference for any of the peptides (Figure 5b). To see if this was due to the shortage of reference TCRs, we sampled 1,000 sequences per pMHC and computed F1@1000 on the limited target sequences. This time, for the FTDALGIDEY peptide, TCRT5 correctly sampled 1 out of 12 sequences at rank 514. To understand if this was better or worse than random sequence generation, we calculated sequence recovery rates for TCRT5 and soNNia-generated sequences. TCRT5 achieved a higher mean sequence recovery rate across all peptides (Figure 5c).

**Figure 5:**
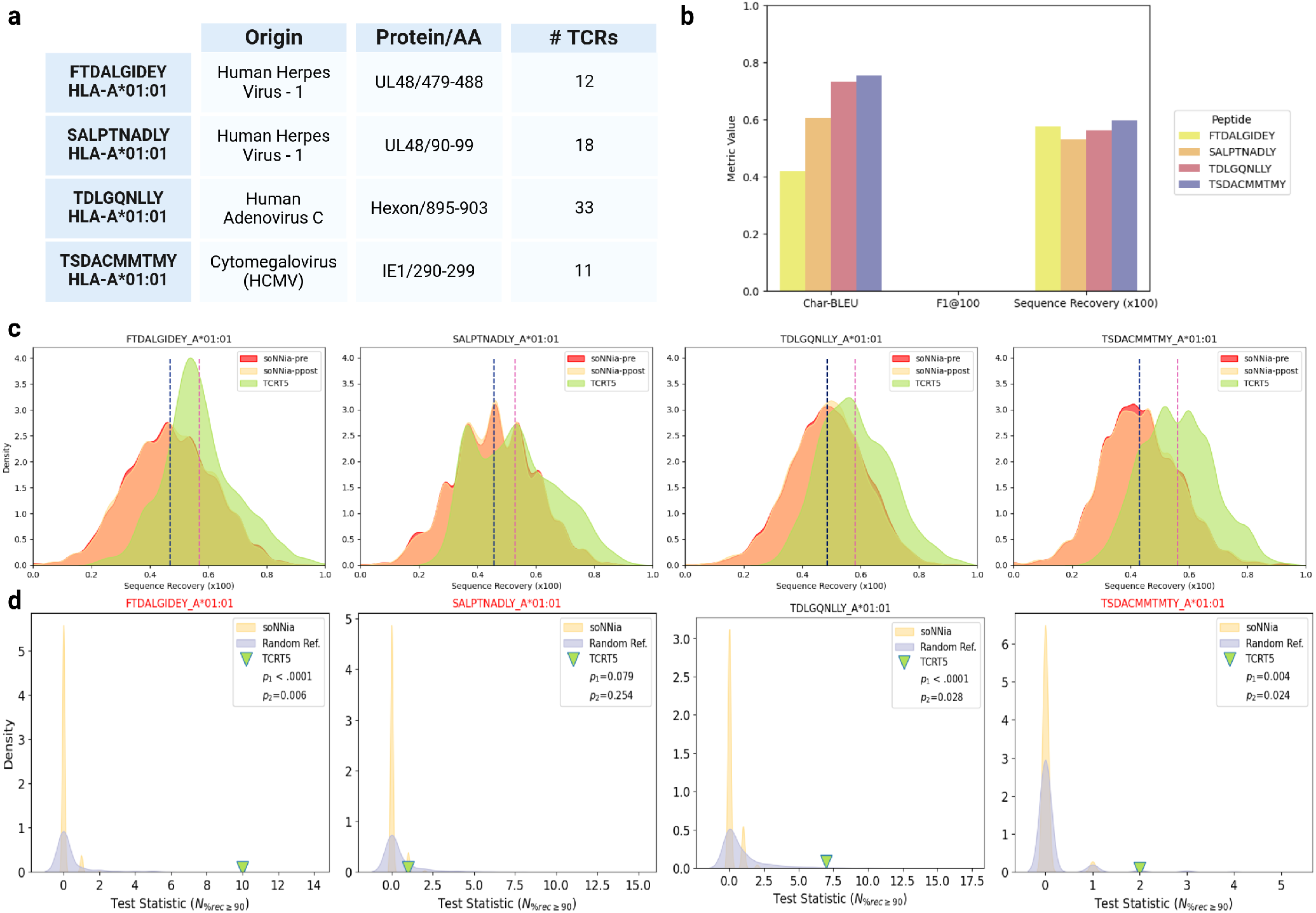
TCRT5 Performance on IMMREP23 “Unseen” Epitopes. (a) Table of unseen pMHCs along with their species, originating proteins’ amino acids, and the number of validated cognate TCRs. FTDALGIDEY, SALPTNADLY, TSDACMMTMY are designated “unseen” epitopes from IMMREP23 since they do not appear in IEDB and VDJdb. Here we include TDLGQNLLY in the unseen epitopes given its absence from our training set. (b) Accuracy metrics colored by epitope for 100 generations. Mean average precision is omitted given an F1 score of 0. (c) Distribution of sequence recoveries are shown for 1000 conditional generations from TCRT5 against 10000 generations from the unconditional soNNia `‘-ppost’` and `‘-pre’` generative models. (d) Bootstrapped test of significance. Our chosen test statistic is the number of generations that have at least a 90% sequence recovery rate for a fixed number of generations (*k*), in this case *k*=1000. We compare the observed test statistic using TCRT5’s generations against the soNNia (yellow) and random reference (purple) null distributions. The soNNia null is constructed by sampling 1000 sequences from soNNia `‘ppost’` and computing the sequence recoveries against the ground truth CDR3*β* sequences. The random reference null uses soNNia `‘ppost’` to generate fake ground truth CDR3*β* sequences, equal in number to the ground truth TCRs per epitope, to use for sequence recovery calculations with the TCRT5 generations. Empirical p-values *p*_1_ and *p*_2_ are calculated as the number of cases from null distribution 1 and 2 respectively where the test statistics are greater than or equal to the observed statistic divided by the number of trials (1000). A p-value of 0.0 is binned as <.0001.

While greater mean sequence recovery of TCRT5 generations serves as a preliminary sanity check for desired model function, we sought to further assay whether TCRT5 generated more useful individual sequences than a random generator. Since we found the dynamic range for nonzero F1 scores to occur where the sequence recovery was greater than 90% (See A.3, Figure S2d-e), we checked to see if TCRT5 generated more sequences with ≥ 90% sequence identity to the true cognate sequences compared to random. To do so, we established two independent, but complementary null distributions. The first was constructed by bootstrapping our test statistic against soNNia generated sequences, counting the number of instances where the soNNia CDR3*β* sequences (out of 1000 sequences per simulation) achieved at least a 90% sequence recovery. The second null was constructed by calculating the sequence recovery rate of TCRT5 generations against a reference size-matched sample of random CDR3*β* sequences generated by soNNia. This null was constructed to evaluate the specificity of TCRT5 to a particular epitope-specific repertoire over a random repertoire. We calculated empirical p-values based on the fraction of simulations where the null distributions matched or exceeded the observed TCRT5 performance with the ground truth sequences. We found that for all of the unseen epitopes except one, TCRT5 generations outperformed both nulls at a statistical significant level of 0.05 (Figure 5d). For the soNNia random generations null, we observe *p*_1_-values of <.0001, .079, <.0001, and .004 for FTDALGIDEY, SALPTNADLY, TDLGQNLLY, and TSDACMMTMY, respectively. For the random references null, we observe *p*_2_-values of .006, .254, .028, and .024. These results indicate that while TCRT5 generates more closer to ground truth sequences for a reduced set of cognate TCRs than random, there is some degree of non-specific generation as as well. Taken together, these results highlight that while TCRT5 outperforms random sequence generation for the task of antigen-specific TCR design on even sparsely validated epitopes, it remains unavoidable that wet lab validation is necessary to adequately determine the level of non-specificity versus unconfirmed reactivity in the generations.

## 4 Discussion

In this study we set out to design real antigen-specific TCR sequences for unseen epitopes and characterize the nature of the generations beyond their metrics. Building on our prior work framing the conditional TCR design problem as a sequence-to-sequence (seq2seq) task, we explored concepts from low-resource machine translation techniques to address the extremely limited labeled data in the TCR-epitope specificity domain. We introduce novel approaches in this context including data augmentation techniques, joint pre-training, and bi-directional sequence generation to help maximize performance on both accuracy and diversity metrics. We then evaluated our models against well-studied pMHCs to understand model behavior before selecting an optimal model and applying it on real world scenarios.

Working around a need to hold out dense evaluation data for the improbable likelihood of exact sequence recovery, we constructed a target-rich validation set, sacrificing the multiplicity of their TCR:pMHC mappings from our training dataset. In order to re-introduce some depth into our training data, we injected a large volume of semi-synthetic examples drawn from the MIRA [59] dataset, skewing the parallel dataset towards a majority SARS-CoV2 TCR sequences. Remarkably, the models demonstrated an ability to generalize outside the SARS-CoV2 specific repertoire, achieving exact sequence matches to other reference CDR3*β* sequences associated with viral epitopes and neoantigens. This level of retained input sensitivity, while not perfect, suggests robustness to data imbalance for the, which can help inform *in-vitro* data generation strategies on depth vs. breadth-first discovery of antigen-specific TCR for rare pMHCs.

After establishing the superiority of our baseline models over unconditional generation, we trained ten additional model variants on additional tasks, leveraging denoising and bi-directional training to learn the intra- and inter-sequence co-dependencies that exist in and between the TCR and pMHC. Surprisingly we found that pre-training had opposite effects for TCRBART which was pre-trained on BERT’s token masking objective, and TCRT5 which was trained on the original T5 span masking. Since both models were pre-trained using a masking rate of 15%, we suspect that the higher order k-mers learned by span masking are better suited for learning CDR3*β* sequences, though more work is necessary to determine exactly what is driving these changes and if a different set of hyperparameters improves TCRBART’s performance. Between TCRBART and TCRT5, the bidirectional and multi-task models achieved higher sequence recovery and an increased number of non-zero F1 scores across validation pMHCs, indicating that these strategies may be beneficial for capturing antigen-specific motifs. However, upon closer examination of the actual sequences, we discovered that these multi-task models also had an increased propensity to sample polyspecific TCRs, as perceived by the models, that bind numerous antigens and exploited their prevalence in the validation set. We suspect that this may be due to the target collapse going in the reverse TCR → pMHC direction wherein a large number of diverse TCRs collapse onto the few well-studied and well-represented training set pMHCs, and from them the high abundance TCRs are re-sampled in a self-consistent manner [49]. In evaluating model performance, we observed that it correlated with the density of reference data for each pMHC. Notably, between metrics, we found a significant increase in F1 sensitivity above a 90% sequence identity threshold, suggesting this metric may be a more meaningful indicator than mean sequence recovery for assessing antigen-specific repertoire quality.

Taking these lessons into account, we found TCRT5 to be the most well-rounded model for its high accuracy, diversity, and attenuated reliance on polyspecific generations. We rigorously benchmarked it, highlighting a reduction in sequence diversity driven by preferential sampling of sequences with high V(D)J generation probabilities via beam search. Still, we show TCRT5’s utility by generating validated antigen-specific CDR3*β* sequences not encountered during training. We further demonstrate its greater than random performance on the IMMREP2023 “unseen” antigens, possessing substantially fewer known cognate TCRs, providing a pathway for generating functionally relevant TCR repertoires for sparsely sampled epitopes likely to be encountered in a real therapeutic scenario.

### 4.1 Limitations

The current iteration of our study has many limitations, stemming from both our approach and an innate paucity of available data. First, is our reduction in the sequence complexity to focus only on the CDR3*β* loop of the TCR, even though the *α* chain and other determinants of TCR identity such as the V and J gene have been shown to play an important role in determining specificity [27, 87]. In its current state, our model requires template TCRs for which the CDR3*β* can be designed.

Second is the limitation of our sequence-based evaluation approach, which requires at least one known target TCR sequence to which our sequence based metrics can be computed (F1, Char-BLEU, sequence recovery). Thus, our evaluation framework is not extensible to evaluate model performance on novel epitopes for which no known TCRs have been discovered. Nevertheless, we strongly believe that recapitulation of known binders to a target-rich validation is required, given a still-developing understanding of T cell activation working to reconcile suboptimal structural and biochemical features with potent responses. As our understanding and ability to predict TCR recognition improve, this highly limiting metric may be loosened. However, in the interim, we advocate for its inclusion as as a necessary but not sufficient standard to establish successful training before interrogating other proxies for real-world utility.

Last, given the relatively sparse nature of data in even the largest epitope-specific repertoires, we expose ourselves to a high degree variance across all model performance metrics when focusing on sequence sequence-based metrics that center around a reference set. To mitigate this variance, we leaned on our target-rich dataset to not only choose the best model/checkpoint but also study the models’ behavior under a strict success criterion. We understand and accept that this introduces leakage through model selection. However, we argue for its necessity given the severe data sparsity to evaluate pMHCs stemming from multiple disease contexts. Importantly, TCRT5 demonstrates consistent, monotonic improvement in performance across training checkpoints, suggesting that our selection reflects real learning rather than arbitrary fluctuations in performance driven by random permutations to a model’s hyperparameters. Additionally, after using this split to understand these models’ behavior on unseen epitopes (under an optimal checkpoint), we evaluate our lead model on the IMMREP epitopes not used for training or in validation.

### 4.2 Future Directions

While our study set out to understand and improve upon the potential of generative models to design antigen-specific TCR sequences, several exciting areas remain for further exploration and improvement on this front. Given the marked reduction in sequence diversity, exploring novel decoding strategies may help address the bias towards high likelihood TCRs. Techniques from reinforcement learning that promote diversity and other methods of refining model outputs such as steering could also help increase performance by principled sampling of diverse sequence motifs. On the model architecture front, it is unlikely that any major modifications will greatly increase performance on our metrics, under the current data sparsity. Thus it is essential to conduct extensive *in-vitro* validation of model-generated CDR3*β* sequences. Mapping the correlation between various metrics to functional validity will be of great significance to the field in assessing generations, especially in the novel epitope setting. In addition, experimental validation will also be critical in revealing to what degree the model polyspecific generations are noise and the cases in which the polyspecific sequences are actually antigen-specific. Finally, in order to realize the potential of sequence-to-sequence models as an accelerant to therapeutic agents, more work must be done to extend these methods paired and full chain TCR representations so that true de novo TCRs can be generated expressed *in-situ*.

### 4.3 Conclusion

One of the most fundamental questions in immunology is how T-cell receptors (TCRs) achieve precise specificity for peptide-MHC (pMHC) complexes derived from atypical cellular contexts, enabling them to distinguish between self and nonself peptides with incredible sensitivity. This specificity, however, is inherently paradoxical, as TCRs must recognize a pathogen space that prohibits 1:1 mapping of TCRs and pMHCs, given resource constraints. This complex many-to-many relationship between TCRs and pMHCs has posed a significant challenge to modeling efforts. Addressing this challenge would greatly advance our understanding of T cell biology, offering new insights into the principles governing adaptive immunity receptor specificity and providing a foundation for diverse applications in cutting edge therapeutic modalities.

Our work demonstrates the potential of generative models to address a key challenge in TCR research: producing antigen-specific repertoires with high fidelity and diversity in a data-scarce domain. By exploring low-resource machine translation techniques and novel training strategies, we highlight the potential and the many remaining steps toward developing methods that can generate TCRs for specific antigens, even in the absence of extensive experimental data. This capability has profound implications for cellular therapies, enabling the rapid generation of TCRs that can be screened for cross-reactivity with self-epitopes, thereby streamlining the traditional TCR discovery process, which typically relies on a resource-intensive *in-vitro* discovery pipeline. While our study reveals the benefits and limitations of these models in capturing TCR specificity, it also underscores the need for further validation to bridge the gap between computational predictions and functional relevance. As more data becomes available, both model performance and evaluation metrics are expected to improve, moving the field closer to scalable, high-precision TCR design for precision immunotherapy.

## A APPENDIX

## A.1 Data and Code Availability

All sequence data (source-target pairs), computed results, and code used for training and evaluating TCRT5 can be found on: https://github.com/pirl-unc/tcr_translate. For ease of use, the model and tokenizer for TCRT5 can be found on HuggingFace at: https://huggingface.co/dkarthikeyan1/tcrt5_ft_tcrdb. Additionally, the pre-trained TCRT5 may be found at https://huggingface.co/dkarthikeyan1/tcrt5_pre_tcrdb.

## A.2 Model Architecture

We choose BART [43] and T5 [44], both encoder:decoder transformer models, for the model class’s demonstrated performance on robust benchmarks spanning various seq2seq tasks in the broad NLP setting [88, 89]. Compared to the encoder-only transformers like BERT [90], which are great at learning rich feature representations of sequences, and decoder-only models such as GPT[91] that are adept at generating coherent text from prompts, the encoder-decoder architecture combines learning rich latent representations of input sequences and sampling meaningful target sequences in a robust manner. This is especially useful when the mapping between source and target sequences is of different lengths, different languages, or otherwise complicated [92]. Both TCRBART and TCRT5 are thus specific implementations of bi-directional encoders coupled with autoregressive decoders, whose transformer-block architectures do not diverge from their inspirations (Additional details may be found in the original papers). Given the many-to-many mapping of the problem and somewhat varying lengths between sequences, we leverage the encoder-decoder framework to learn rich representations of the pMHC which is used to sample diverse and accurate CDR3*β* sequences by the decoder. There are a few differences in the actual implementations of the original BART and T5 models. While the BART model follows the implementation of the original encoder:decoder transformer introduced in [93], T5 makes a few notable changes (Figure 1 b). The main differences are in the implementation of the layer norm (T5 implements the root mean squared LN and does away with a linear bias), the location of the layer norm (T5 opts for LN prior to the attention mechanism), the encoding of positional information (T5 uses a linear attention bias to encode relative positional information), and dropout (T5 uses dropout gratuitously across the flow of information through the model). The intuition and effects of these modifications can be found in the original T5 paper [44].

## A.3 Metric Correlation

When we expanded out the twelve models’ performance by pMHC, we noticed that the models’ performance was largely conserved, indicating that some pMHCs were predisposed to higher performance than others (Figure S3a-b). We investigated the correlation of two of our performance metrics (F1@100 and sequence recovery) against features of our validation examples to see if we could find any associations between performance and characteristics of the data, focusing on features that we felt had the potential to influence performance: the similarity to a training set pMHC, the target overlap with the closest known pMHC, and the number of known binders to the validation pMHC (Figure S3c). Of these comparisons, we found that as the number of references increased, the sequence recovery scaled accordingly (Spearman’s *ρ*=0.69). F1, on the other hand had a slightly noisier relationship with the potential explanatory features with target set size being the strongest correlate (Spearman’s *ρ*=0.58). In general, we observed that the examples with a larger number of reference target sequences had a greater sensitivity to a nonzero F1 score. We then proceeded to check the correlation between F1 score and the other metrics (Figure S3d). Of these, we observed a dynamic range of F1s when the mean sequence recovery for an input reached 90%. To test if this was true on an individual sequence level, we evaluated the sequence recovery of all of the translations across all of the models after taking out exact matches from the reference set and saw if there was any separation between the sequence recoveries of known binders and those of unknown specificity. To our surprise, the threshold of 90% persisted and we saw a strong separation where known binders shared at least a 90% to another CDR3*β* sequence in the reference target set for a particular pMHC (Figure S3e). We posit that the 90% sequence identity to known binders may serve as a potentially useful proxy for F1 score in cases where few target sequences exist for an input pMHC and note a very similar finding on a similar edit distance variability around certain CDR3*β* and CDR3*α* motifs for a particular epitopes [94].

## A.4 Quantitative Break Down of Data Constraints on Model Performance

For all metrics, Seq2Seq performance is predicated on a representative number of reference target sequences. Given that we are evaluating generations where even the most represented pMHC has on the order of ≈ 10,000 observed sequences out of a theoretical max of 10^6^ [95, 16] (1% of the total diversity), we are in a regime where model performance is the lower bound on performance. This is because we are evaluating the model not only on its ability to generate correct target sequences, but are inadvertently asking it to generate target sequences that resemble those that have been experimentally validated, completely unrelated to sequence veracity (Figure S1a). We distill this intuition into a probabilistic framework to contextualize the limits on recall-based metrics (i.e. F1 score) for model evaluation, given the amount of data that currently exists:

Assume we have a held-out pMHC (*pMHC*_*i*_) with a theoretical set *C* of cognate CDR3*β* sequences, of which a subset of sequences have been experimentally validated (observed). We can model the likelihood of the generated sequences belonging to the observed set using a composite distribution linking the binomial and hypergeometric distributions. Given model that samples *n* CDR3*β* sequences conditioned on that pMHC, we can define Z to be an unobservable binomially distributed random variable representing the number of correct (but not necessarily observed) generated CDR3*β* sequences.

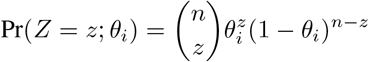

where: *Z* = Random Variable: unobservable number of correct sequences that are in the reference set

*n* = Number of generated translations

*θ*_*i*_ = True model accuracy for a given *pMHC*_*i*_

Then we can construct a conditional distribution of number of correct and observed sequences *Y* |*Z* according to:

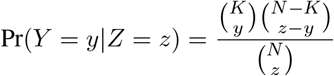

where: *Y* = Random Variable: observed number of correct sequences that have been experimentally validated.

*N* = Number of total cognate sequences (ground truth, partially observed)

*K* = Number of experimentally validated cognate sequences

*n* = Number of generated sequences

*z* = Sample size (number of correct generated sequences, observed through Y)

This gives a joint distribution:

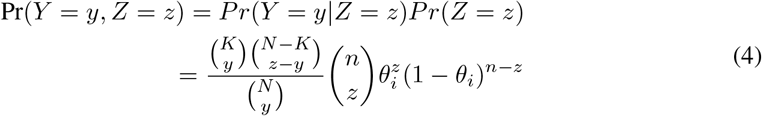

By marginalizing on Y we get the following equation, whose PMF we plot in Figure S1b:

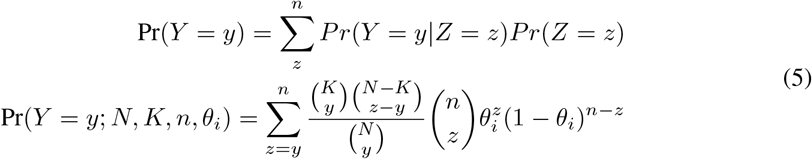

We posit that this framework may be useful in characterizing model performance via estimating the parameter *θ*_*i*_ for pMHCs using Bayesian methods or jointly estimating *Z*_*i*_ and *θ*_*i*_ through the Expectation Maximization algorithm. We leave this for future exploration and use the above for contextualizing evaluation performance in the current data regime.

## B Supplementary Methods

### B.1 Dataset Construction

First, to aggregate the data spanning various sources, formats, and nomenclature, we mapped the columns from each individual dataset to a common consensus schema and concatenated the data along the consensus columns. In the interest of data retention, missing values were reasonably imputed according to other information for that data instance. To keep only the cytotoxic (CD8+) T cells, we filtered the instances wherever the cell-type was provided or where the HLA-Allele was of MHC-class I. In cases where an HLA-haplotype was provided instead of the specific HLA-allele, as was the case for the MIRA data, we used MHCFlurry2.0 [60] to predict the best presenting allele for the given epitope among the potential options from the haplotype information. This data augmentation step resulted in a 5-fold expansion of our training data. An key caveat to note is that the additional training examples are all derived from a single disease context (SARS-CoV2), severely skewing the parallel data’s distribution. Additionally there is room for slight error given the peptide-MHC assignment in-silico and not validated experimentally. However, given the merits of including examples with thousands of TCRs against an epitope, we argue for its inclusion. Additionally, during prototyping, we found its inclusion to increase performance across multiple metrics. Where the granularity of the HLA-information or TR genes was at the serotype level, we inferred the canonical gene/allele by starting off with the subgroup ‘*01’ and incremented it until a matching IMGT gene was found. This step has the potential of introducing minor differences between the unknown ground truth and the imputed pseudo-sequence, as the pseudosequence is well conserved within serotype. Once the data was aggregated and values were imputed, we applied the following column-level standardization for each source of information:

- **Complementarity Determining Region (CDR3***β***), Epitope, and MHC Pseudo-Sequence**: All amino acid representations were normalized used the `‘tidytcells.aa.standardise’` function found in the TidyTcells python package [96].
- **TR Genes**: The TidyTcells package [96] was once again used to standardize the nomenclature surrounding the T-Cell Receptor genes (e.g. TRB-V and TRB-J).
- **HLA-Allele**: HLA alleles were imputed where allele level information when necessary and then normalized using the MHCgnomes package to the standard HLA-[A,B,C]*XX:YY format.

Finally, given the importance of training LLMs on non-redundant data [61] a de-duplication step was performed to consolidate training examples where examples pMHCs that bound to the same CDR3b and shared a k-mer overlap of at least 6 were clustered together and a single representative pair was chosen from each cluster. The allocation of sequence pairs from each cluster was done to balance pMHC representation.

### B.2 Hyperparameter Optimization

Though both the BART and T5 models come with off the shelf recommendations for configurations that are empirically well suited for a number of NLP tasks, we sought to find the set of parameters best suited for the reduced vocabulary size and more tightly bounded sequence lengths when compared to natural language tasks. The impact of scaling in data constrained settings is marginal as shown by recent studies [97] that demonstrate the interplay between the data and model size. Even in this well defined setting, the number of parameters to tune is still considerably large so we use a grid-based sweep to determine an optimized set of hyperparameters for the BART and T5 architectures for our case.

We first investigated the impact of course grained choices in the larger model such as the width and depth by sweeping over model architecture and training algorithm values. For the model architecture we varied the number of attention heads, batch size, *d*_*model*_, feed forward layer dimension, and number of total layers. All models were trained using the cross entropy loss with the AdamW optimizer. For the optimizer, we varied the learning rate and weight decay parameters. To compare model parameters from both the zero-pre-training and pre-training+finetuning regimes, we ran a sweep on samples of the ‘monolingual’ (unlabeled TCR and pMHCs) and ‘parallel’ (paired TCR:pMHC) corpuses to tune performance on the pre-training and seq2seq task, respectively. Due to time and compute constraints, the sweeps were performed on a reduced sample of of 100k TCRs and 100k pMHCs from the monolingual texts for pre-training. Since the parallel corpus was many orders of magnitude smaller to begin with, we took all the pairs not derived from the MIRA dataset (100k paired TCR:pMHC examples), as no allele imputation was used. Interestingly, we found that while the optimal configurations for the BART models were relatively consistent, the optimal T5 configurations for pre-training was a deeper, more narrow network and the one for direct training was a wider more shallow network. To reconcile these differences for a single, task-agnostic, set of optimal parameters, we adopted the following heuristic: if the values were close, the one that resulted in an overall higher parameter count was chosen whereas if the values were far, an intermediate value was chosen. Slight adjustments were performed at the layer count level to adjust for parameters and make TCRBART and TCRT5 comparable. These values are reported in (Table S1). Surprisingly, we found that smaller models yielded better performance, deviating from the hyper-parameters used in our previous works as well as the original BART and T5 papers. The final TCRBART architecture uses 6 encoder and decoder layers at *d*_*model*_ = 768, totaling around 46 million parameters. The final TCRT5 implementation used *d*_*model*_ = 256 and 10 encoder and decoder layers for a total of 42M parameters.

### B.3 Checkpoint Selection

In deciding to choose which checkpoints to use for each model, we observed a marked difference between the performance dynamics of the models with and without pre-training. This is most clearly observed when plotting the diversity and accuracy metrics for each of the model checkpoints, showing distinct training trajectories in the utility space (Figure S3a-b). The pre-trained models demonstrate asymptotically increasing model performance across checkpoints for both the F1@100 and native sequence recovery metrics while the models that were trained directly from random initialization showed signs of potential overfitting, as both metrics peaked early on during training and dropped over additional iterations (Figure S3c-d). This was observed on reduced learning rates as well, indicating a possible regularization effect from pre-training[98]. A distinct difference between TCRBART and TCRT5 variants was the effect of including pre-training. While TCRT5 showed a significant improvement given pre-training, TCRBART showed worse performance. However, the finetuning’s performance dynamics proved to be more stable than the non-pre-trained version, complicating the benefit of adding pre-training to TCRBART. Additionally we examined the number of unique sequences for the models over the checkpoints and saw that the TCRBART-0 and TCRT5-FT showed increasing number of unique sequences over training steps (Figure S3e). For each of the models, the checkpoint with the best performance on F1 was chosen to characterize the optimal performance from each model variant and assess their comparative advantages.

## C Supplemental Figures

## D Supplementary Tables

## Acknowledgments and Disclosure of Funding

This work would not have been possible without numerous fruitful conversations. We thank Dr. Adam Palmer and Dr. Amy Pomeroy for their suggestions and feedback on data communication and our figures. We thank Dr. Will Valdar for his feedback on our statistical methods. We thank Andy Lee, Sara Peterson, and Julia Webb for lending their creativity, expertise, and help in addition to proofreading our manuscript. Finally, we are grateful to the numerous friends and reviewers from various conference venues including: NuerIPs GenBio, AIRR-C VII, and ICML AccMLBio, for their generous and valuable feedback, whose suggestions helped strengthen many portions of our study.

This work was supported largely by the National Science Foundation Graduate Research Fellowship. The authors have no competing interests to declare.

---

**Supplementary Figure 1:**
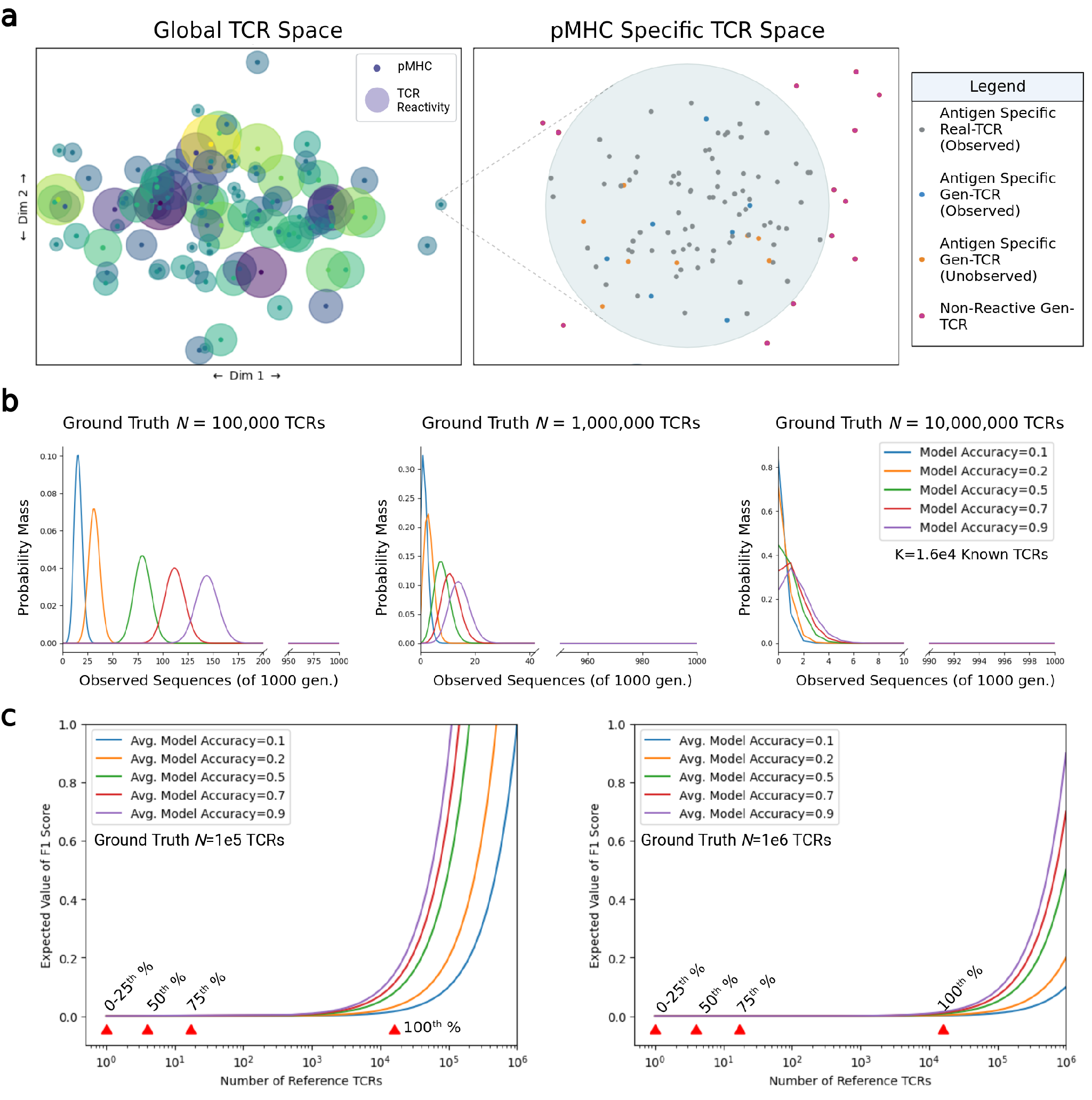
Characterization of the TCR:pMHC Data Landscape. (a) Illustrative diagram showing an idealized rendering of global and local TCR:pMHC context. Global TCR space is represented by 2D projections of various TCR sequences, where antigen-specific TCRs are radially distributed about a pMHC and overlap designates TCR cross reactivity. The local pMHC-specific TCR space shows the distinction between observed TCRs and ground truth TCRs in relation to model generations. (b) Probability mass function (PMF) of Equation plotted for different values of ground truth TCRs and different model accuracies. *K* is fixed to be the current maximum of known TCRs for a given antigen ≈ 16,000. (c) Expected value of the F1 score is plotted for different average model accuracies given *θ* (the PMF simplifies to the conditional hypergeometric distribution). Red arrows indicate the percentiles of reference TCR counts (*K*) from the real data

**Supplementary Figure 2:**
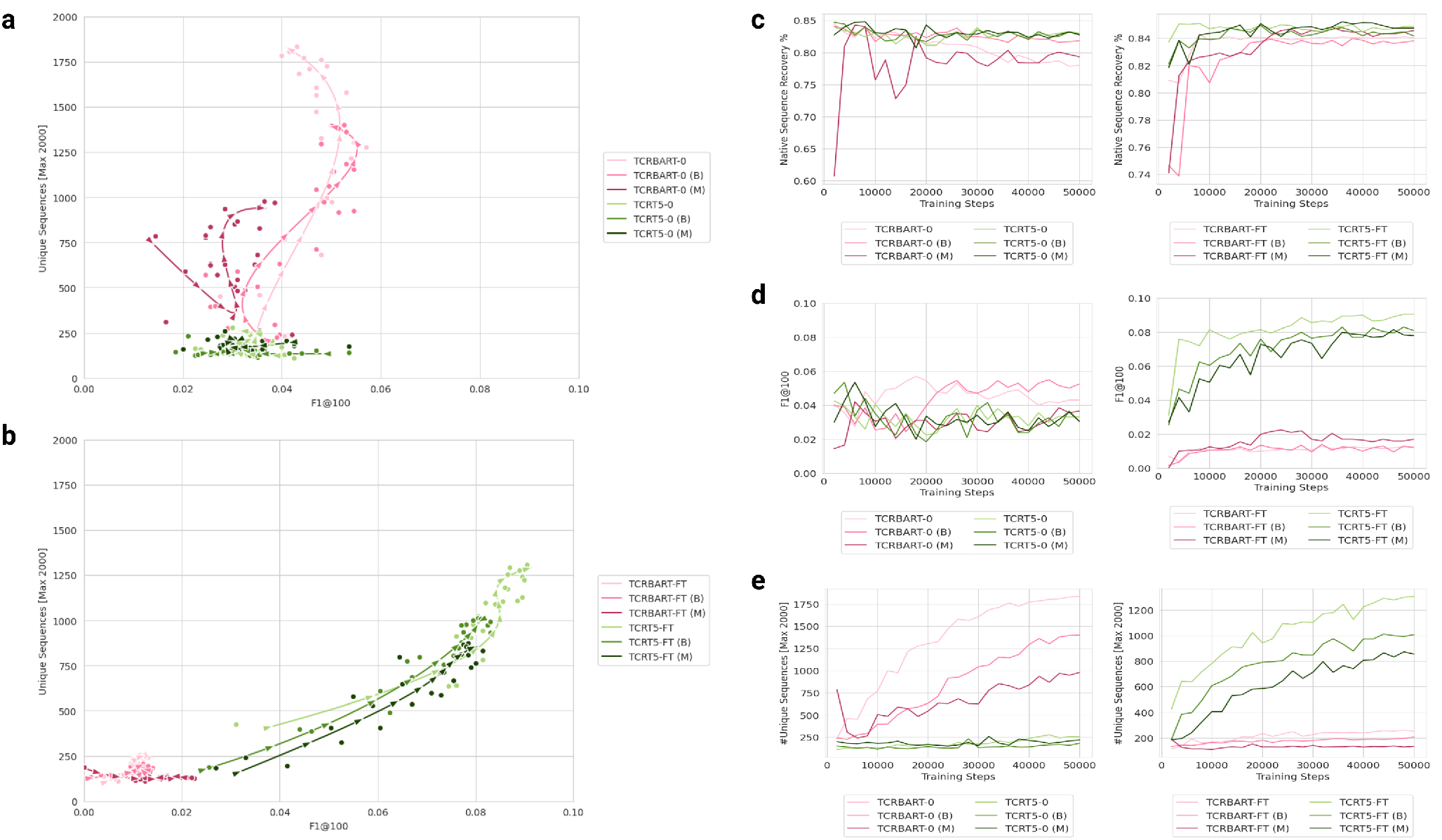
Training dynamics highlight the robustness of pretrained models across checkpoints. Diversity vs. accuracy (F1) plotted for model checkpoints with smoothed interpolated splines and associated arrows showing the direction of model checkpoints through their training trajectory for: (a) Randomly initialized models (zero pre-training) (b) Pre-trained and finetuned models (c) Native sequence recovery for each checkpoint, colored by model, with panel split by pre-training status. (d) F1@100 for all checkpoints by pretraining status. (e) Number of unique generations for each checkpoint across training for all models with panel split by pre-training status. All models checkpoints were taken every 2000 steps across 20 epochs.

**Supplementary Figure 3:**
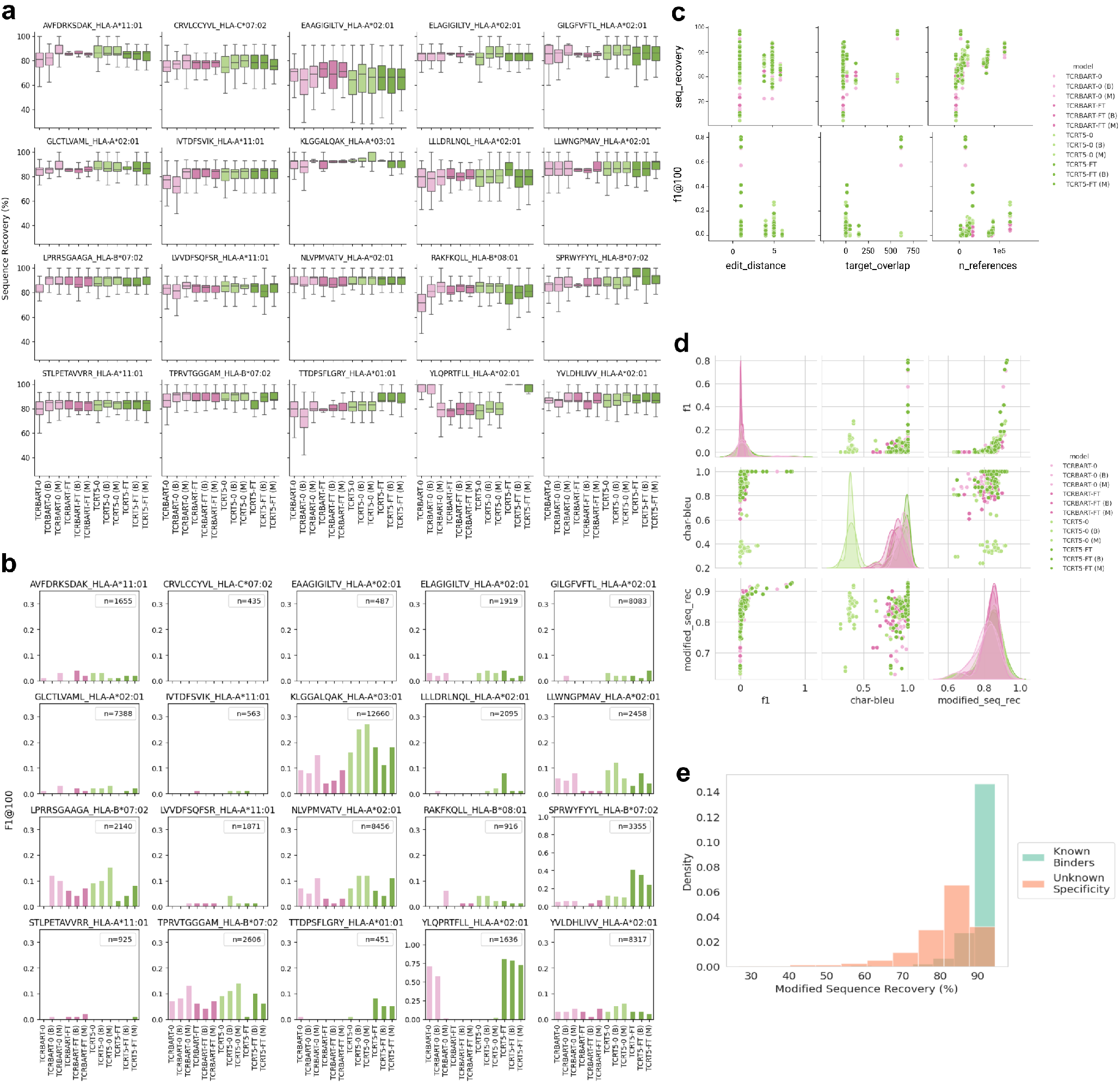
Atomic Metrics. (a) Box and whisker plot of sequence recoveries split by individual pMHC and model. (b) Barplot showing F1@100 score per model and pMHC. Each subplot is demarcated with the number of reference CDR3*β*s in the top right corner. (c) Scatterplot showing relationship between accuracy metrics (sequence recovery and F1@100) and input features (edit distance to closest training pMHC, TCR overlap with closest training pMHC (by edit distance), and number of references (known TCR binders). (d) Correlation plot between sequence-derived metrics. Pairplot showing the pairwise relationships of F1@100, Char-BLEU, and (modified) sequence recovery, across model variants. Modified sequence recovery is calculated by first removing exact matches to the generated sequences from the reference sets and calculating sequence recovery to the closest sequence. (e) Histogram of modified sequence recovery values stratified by known binding status.

**Supplementary Figure 4:**
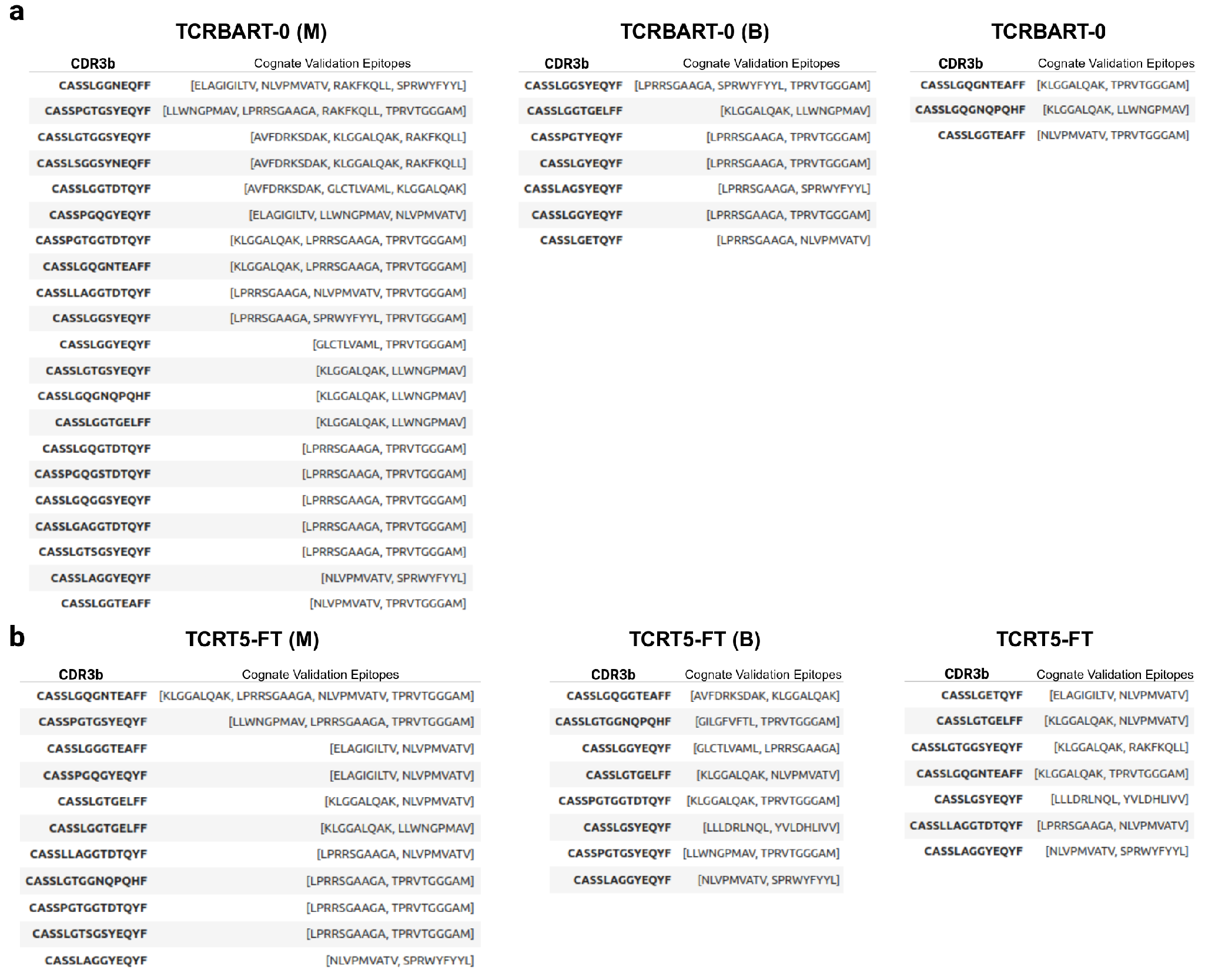
Multi-task models sample more known validated polyspecific TCR sequences. (a) Subset of TCRBART-0 generations across model variants that are known binders to more than one validation pMHC (may be from the same disease context). (b) Subset of TCRT5-FT generations across model variants that are known binders to more than one validation pMHC (may be from the same disease context). Each row is an individual CDR3*β* sequence that was generated for and found in the experimentally validated set of reference TCRs for the listed validation pMHCs.

**Supplementary Figure 5:**
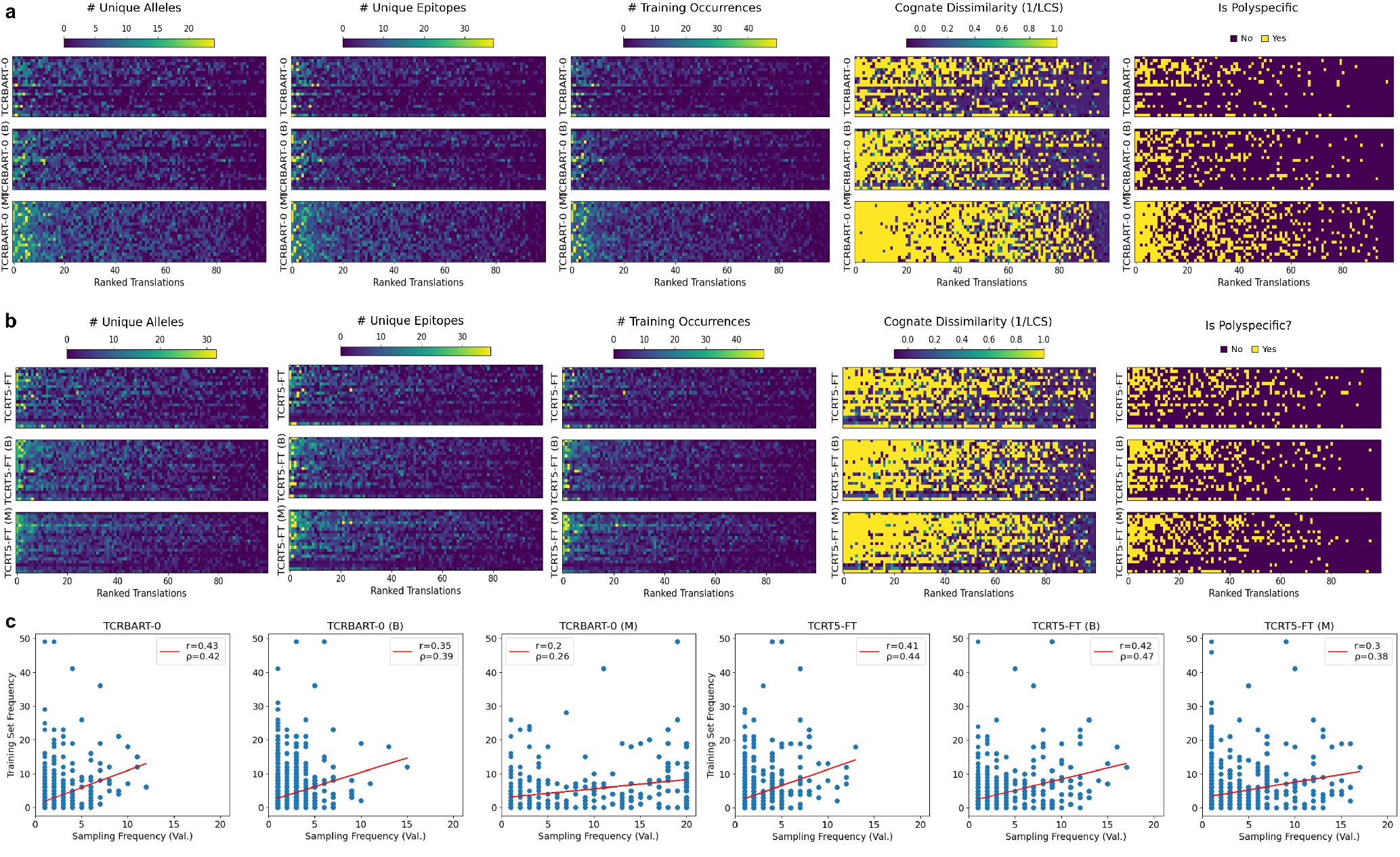
Exploring polyspecificity vs. training set statistics across baseline, bidirectional, and multi-task model variants. (a) Heatmap of ranked TCRBART-0 translations across pMHCs colored by number of known alleles, known epitopes, training set frequency, epitope dissimilarity, and membership status in the 915 polyspecific TCRs. (b) Analogous heatmap as panel ‘a’ but for TCRT5-FT generations. (c) Correlation plots for TCRBART-0 and TCRT5-FT model generations and training set occurrence. Line of best fit is shown in red. Pearson’s *r* and Spearman’s *ρ* are also provided for each model.

**Supplementary Figure 6:**
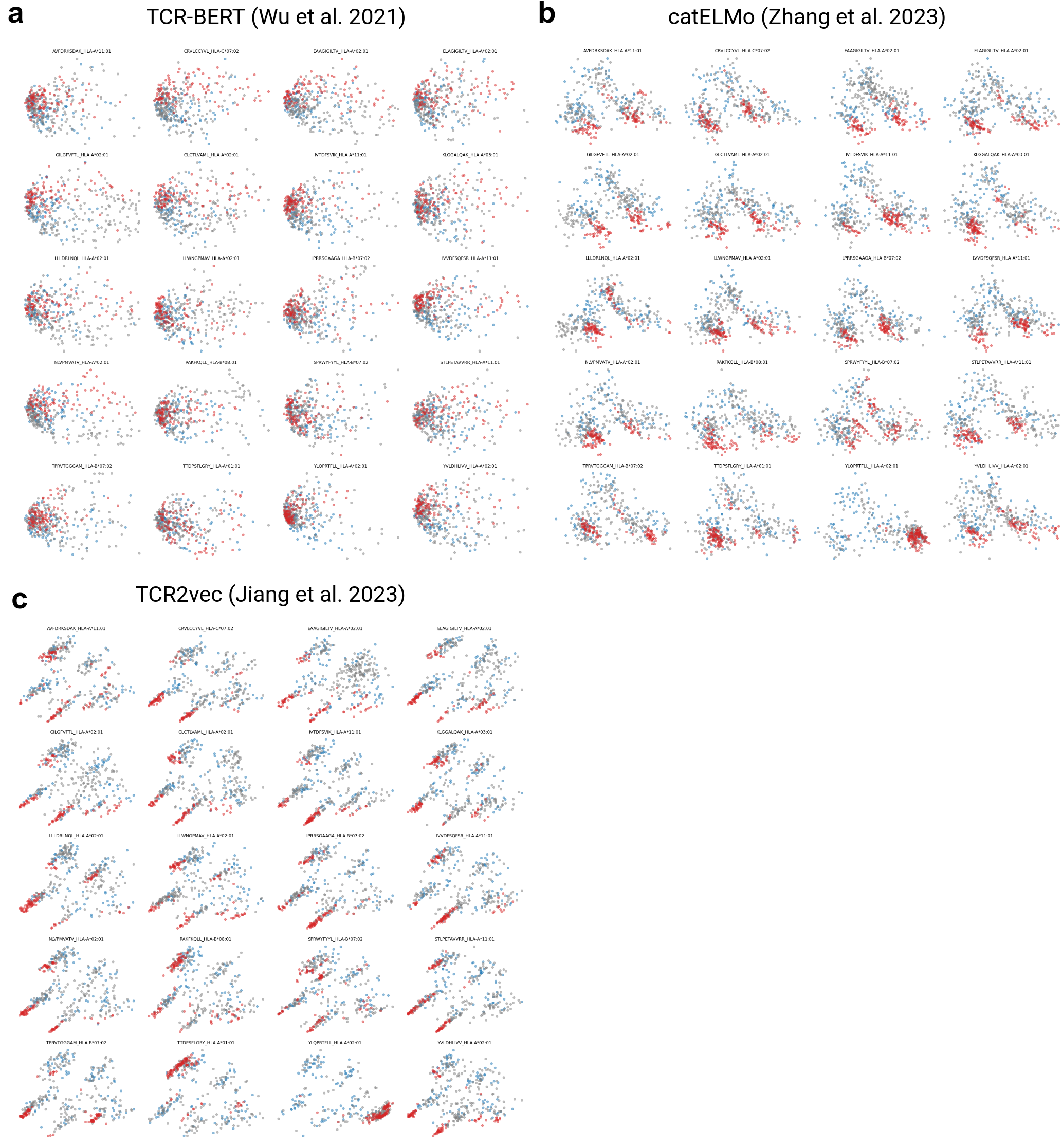
CDR3*β* embeddings highlight reduction in sampled TCR space. PCA dimensionality reduction of embeddings generated by sequence based methods are shown for: (a) TCR-BERT (b) catELMo (c) TCR2vec. Red points indicate sequences generated by TCRT5, gray corresponds to reference translations, and blue points are soNNia generated sequences. Reference TCRs are downsampled to 200 sequences and 100 background sequences are shown.

**Supplementary Figure 7:**
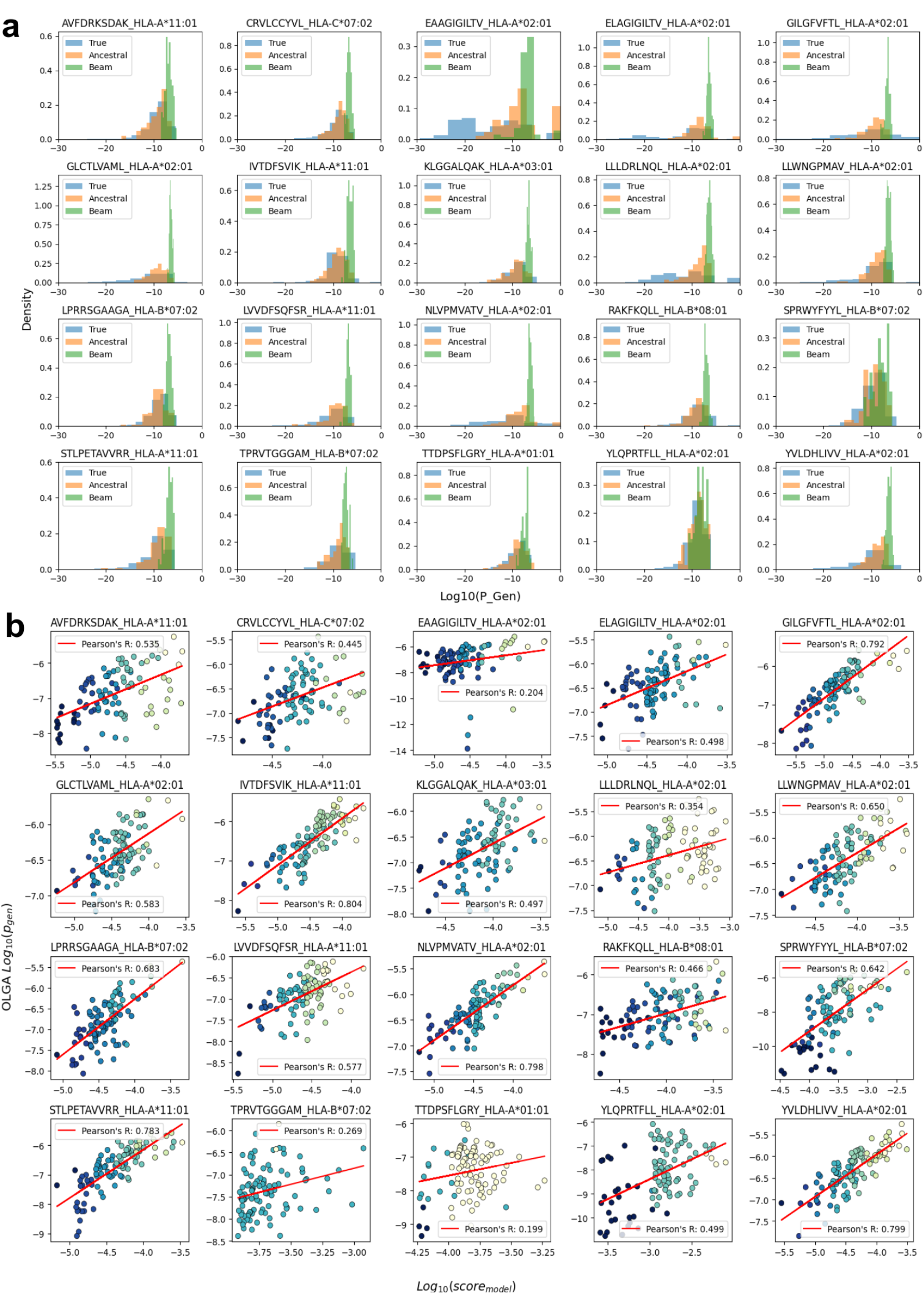
TCRT5 Sequence Likelihoods. (a) Histograms showing the OLGA *p*_*gen*_ values for the reference CDR3*β*s as well as those generated by beam search and ancestral sampling methods. (b) Correlation plots showing the model scores (model sequence likelihoods) against the biophysical OLGA *p*_*gen*_. Axes are log10 scaled. Red line is the best fit line with associated Pearson’s R.

**Supplementary Figure 8:**
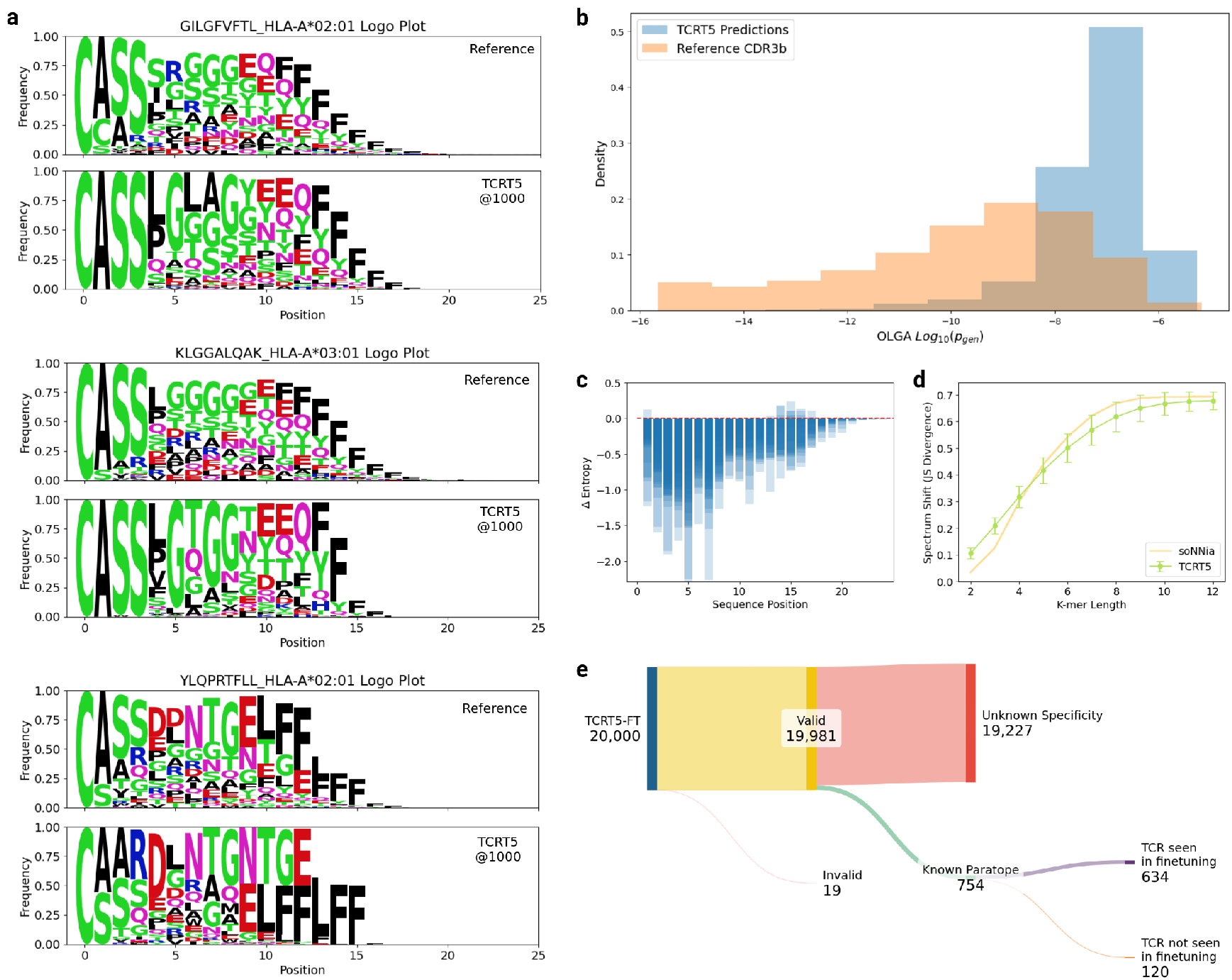
TCRT5 Metrics @1000. (a) Sequence logo plot generated from TCRT5 for the canonical GILGFVFTL (Influenza A), KLGGALQAK (CMV), and YLQPRTFLL (SARS-CoV2) from 1000 generations instead of 100. (b) TCRT5@1000 with beam search still preferentially samples sequences at the right tail of OLGA generation probabilities. (c) Generated sequences experience a decrease in Shannon entropy across most positions, however, some pMHC examples exhibit an increase in entropy compared to reference sequences. Barplots for individual pMHCs are overlaid on one another. (d) K-mer spectrum shift plot showing the Jensen Shannon divergence between generated and reference sequences for TCRT5@1000. Error bars mark the mean and 1-standard deviation across validation pMHCs. Mean soNNia values are shown per simulated run, with 1000 generations per pMHC per run over 100 simulations. (e) Sankey diagram of TCRT5@1000 generations showing the validity as measured by nonzero generation probability, known binding status, and training set membership.

**Supplementary Table 1:** Model Architecture Hyperparameters

|  | TCRBART | TCRT5 |
| --- | --- | --- |
| Parameters | 46M | 42M |
| $d_{model}$ | 768 | 256 |
| Vocab Size | 28 | 128 |
| Encoder Layers | 6 | 10 |
| Decoder Layers | 6 | 10 |
| Max Position Embedding | 512 | 512 |
| Attention Heads | 16 | 16 |
| Feed Forward Dim | 128 | 1024 |
| Cross Attention | ✓ | ✓ |

**Supplementary Table 2:**
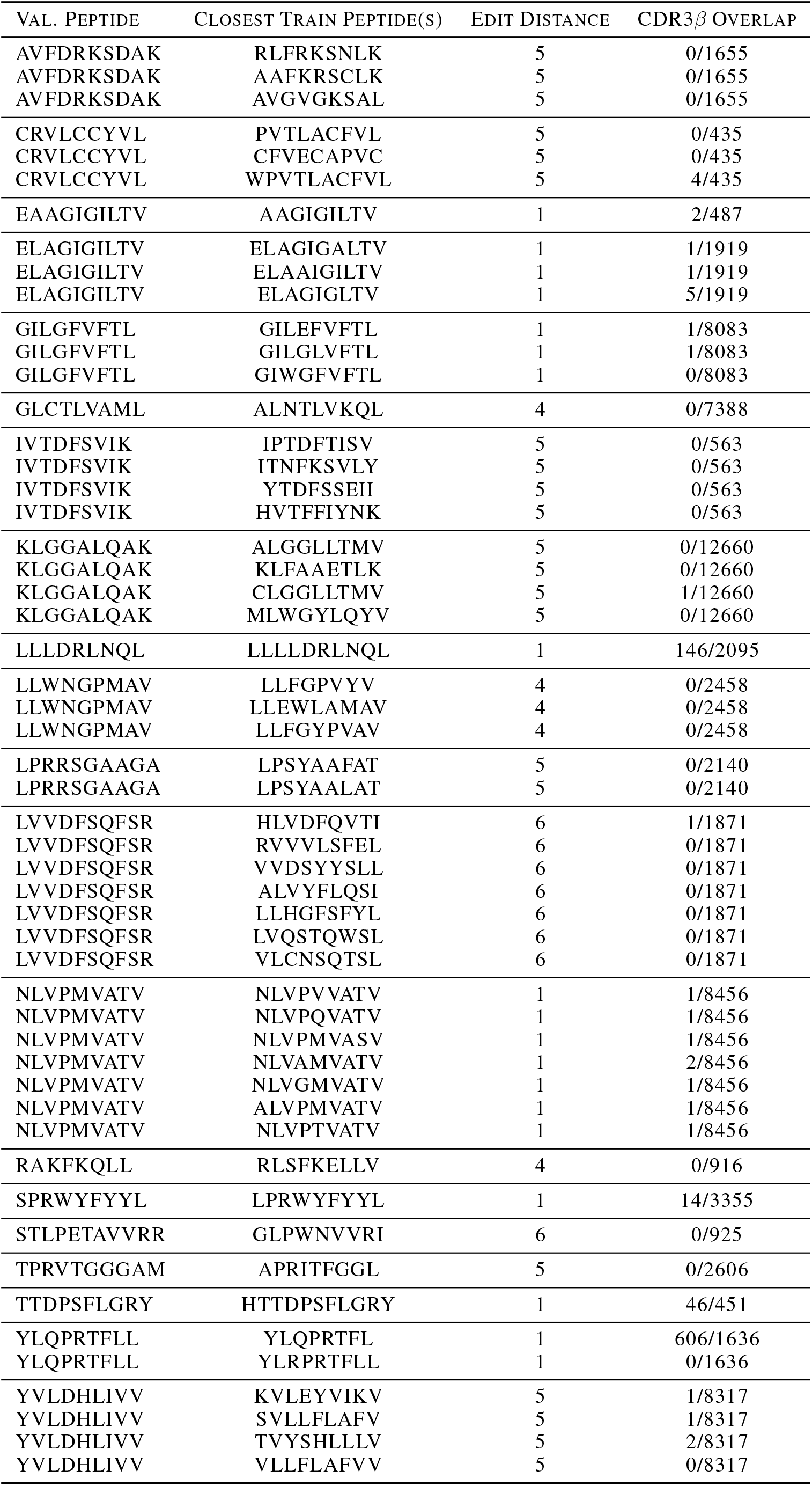
Characterization of Train/Val. Target Overlap

## References

[1] Alexis M. Kalergis, Toshiro Ono, Fuming Wang, Teresa P. DiLorenzo, Shinichiro Honda, and Stanley G. Nathenson. Single Amino Acid Replacements in an Antigenic Peptide Are Sufficient to Alter the TCR Vb Repertoire of the Responding CD8+ Cytotoxic Lymphocyte Population1. The Journal of Immunology, 162(12):7263–7270, 06 1999. ISSN 0022-1767. doi: 10.4049/jimmunol.162.12.7263. URL 10.4049/jimmunol.162.12.7263.

[2] Xingyu Cao, Guangna Liu, Jianping Zhang, Yanli Zhao, Hua Chen, Hongli Zheng, Wei Rui, Lemei Jia, Xueqiang Zhao, Xin Lin, and Peihua Lu. A Novel CMV-Specific TCR-T Cell Therapy Is Effective and Safe for Refractory CMV Infection after Allogeneic Hematopoietic Stem Cell Transplantation. Blood, 138(Supplement 1):3848–3848, 11 2021. ISSN 0006-4971. doi: 10.1182/blood-2021-146446. URL 10.1182/blood-2021-146446.

[3] Ifigeneia Tzannou, Anastasia Papadopoulou, Swati Naik, Kathryn Leung, Caridad A. Martinez, Carlos A. Ramos, George Carrum, Ghadir Sasa, Premal Lulla, Ayumi Watanabe, Manik Kuvalekar, Adrian P. Gee, Meng-Fen Wu, Hao Liu, Bambi J. Grilley, Robert A. Krance, Stephen Gottschalk, Malcolm K. Brenner, Cliona M. Rooney, Helen E. Heslop, Ann M. Leen, and Bilal Omer. Off-the-shelf virus-specific t cells to treat bk virus, human herpesvirus 6, cytomegalovirus, epstein-barr virus, and adenovirus infections after allogeneic hematopoietic stem-cell transplantation. Journal of Clinical Oncology, 35(31):3547–3557, 2017. doi: 10.1200/JCO.2017.73.0655. URL 10.1200/JCO.2017.73.0655. PMID: 28783452.

[4] Yunyu Mao, Qibin Liao, Youwei Zhu, Mingyuan Bi, Jun Zou, Nairong Zheng, Lingyan Zhu, Chen Zhao, Qing Liu, Li Liu, Jun Chen, Ling Gu, Zhuoqun Liu, Xinghao Pan, Ying Xue, Meiqi Feng, Tianlei Ying, Pingyu Zhou, Zhanshuai Wu, Jian Xiao, Renfang Zhang, Jing Leng, Yongtao Sun, Xiaoyan Zhang, and Jianqing Xu. Efficacy and safety of novel multifunctional m10 car-t cells in hiv-1-infected patients: a phase i, multicenter, single-arm, open-label study. Cell Discovery, 10(1):49, May 2024. ISSN 2056-5968. doi: 10.1038/s41421-024-00658-z. URL 10.1038/s41421-024-00658-z.

[5] Christoph T. Ellebrecht, Vijay G. Bhoj, Arben Nace, Eun Jung Choi, Xuming Mao, Michael Jeffrey Cho, Giovanni Di Zenzo, Antonio Lanzavecchia, John T. Seykora, George Cotsarelis, Michael C. Milone, and Aimee S. Payne. Reengineering chimeric antigen receptor t cells for targeted therapy of autoimmune disease. Science, 353(6295):179–184, 2016. doi: 10.1126/science.aaf6756. URL https://www.science.org/doi/abs/10.1126/science.aaf6756.

[6] Fabian Müller, Jule Taubmann, Laura Bucci, Artur Wilhelm, Christina Bergmann, Simon Völkl, Michael Aigner, Tobias Rothe, Ioanna Minopoulou, Carlo Tur, Johannes Knitza, Soraya Kharboutli, Sascha Kretschmann, Ingrid Vasova, Silvia Spoerl, Hannah Reimann, Luis Munoz, Roman G. Gerlach, Simon Schäfer, Ricardo Grieshaber-Bouyer, Anne-Sophie Korganow, Dominique Farge-Bancel, Dimitrios Mougiakakos, Aline Bozec, Thomas Winkler, Gerhard Krönke, Andreas Mackensen, and Georg Schett. Cd19 car t-cell therapy in autoimmune disease — a case series with follow-up. New England Journal of Medicine, 390(8):687–700, 2024. doi: 10.1056/NEJMoa2308917. URL https://www.nejm.org/doi/full/10.1056/NEJMoa2308917.

[7] Rom Leidner, Nelson Sanjuan Silva, Huayu Huang, David Sprott, Chunhong Zheng, Yi-Ping Shih, Amy Leung, Roxanne Payne, Kim Sutcliffe, Julie Cramer, Steven A. Rosenberg, Bernard A. Fox, Walter J. Urba, and Eric Tran. Neoantigen t-cell receptor gene therapy in pancreatic cancer. New England Journal of Medicine, 386(22):2112–2119, 2022. doi: 10.1056/NEJMoa2119662. URL https://www.nejm.org/doi/full/10.1056/NEJMoa2119662.

[8] Charlotte Harrison. Tcr cell therapies vanquish solid tumors — finally. Nature Biotechnology, 42(10):1477–1479, Oct 2024. ISSN 1546-1696. doi: 10.1038/s41587-024-02435-5. URL 10.1038/s41587-024-02435-5.

[9] Andrew Poole, Vijaykumar Karuppiah, Annabelle Hartt, Jaafar N. Haidar, Sylvie Moureau, Tomasz Dobrzycki, Conor Hayes, Christopher Rowley, Jorge Dias, Stephen Harper, Keir Barnbrook, Miriam Hock, Charlotte Coles, Wei Yang, Milos Aleksic, Aimee Bence Lin, Ross Robinson, Joe D. Dukes, Nathaniel Liddy, Marc Van der Kamp, Gregory D. Plowman, Annelise Vuidepot, David K. Cole, Andrew D. Whale, and Chandramouli Chillakuri. Therapeutic high affinity t cell receptor targeting a krasg12d cancer neoantigen. Nature Communications, 13 (1):5333, Sep 2022. ISSN 2041-1723. doi: 10.1038/s41467-022-32811-1. URL 10.1038/s41467-022-32811-1.

[10] Paul Nathan, Jessica C. Hassel, Piotr Rutkowski, Jean-Francois Baurain, Marcus O. Butler, Max Schlaak, Ryan J. Sullivan, Sebastian Ochsenreither, Reinhard Dummer, John M. Kirkwood, Anthony M. Joshua, Joseph J. Sacco, Alexander N. Shoushtari, Marlana Orloff, Josep M. Piulats, Mohammed Milhem, April K.S. Salama, Brendan Curti, Lev Demidov, Lauris Gastaud, Cornelia Mauch, Melinda Yushak, Richard D. Carvajal, Omid Hamid, Shaad E. Abdullah, Chris Holland, Howard Goodall, and Sophie Piperno-Neumann. Overall survival benefit with tebentafusp in metastatic uveal melanoma. New England Journal of Medicine, 385(13):1196–1206, 2021. doi: 10.1056/NEJMoa2103485. URL https://www.nejm.org/doi/full/10.1056/NEJMoa2103485.

[11] Yating Liu, Xin Yan, Fan Zhang, Xiaoxia Zhang, Futian Tang, Zhijian Han, and Yumin Li. Tcr-t immunotherapy: The challenges and solutions. Frontiers in Oncology, 11, 2022. ISSN 2234-943X. doi: 10.3389/fonc.2021.794183. URL https://www.frontiersin.org/articles/10.3389/fonc.2021.794183.

[12] Kazusa Ishii, John S. Davies, Andrew L. Sinkoe, Kilyna A. Nguyen, Scott M. Norberg, Crystal P. McIntosh, Tejas Kadakia, Carylinda Serna, Zachary Rae, Michael C. Kelly, and Christian S. Hinrichs. Multi-tiered approach to detect autoimmune cross-reactivity of therapeutic t cell receptors. Science Advances, 9(30):eadg9845, 2023. doi: 10.1126/sciadv.adg9845. URL https://www.science.org/doi/abs/10.1126/sciadv.adg9845.

[13] Max D. Cooper and Matthew N. Alder. The evolution of adaptive immune systems. Cell, 124(4):815–822, Feb 2006. ISSN 0092-8674. doi: 10.1016/j.cell.2006.02.001. URL 10.1016/j.cell.2006.02.001.

[14] Ekeruche-Makinde Wooldridge, Skowera van den Berg, Tan Miles, Clement Dolton, Price Llewellyn-Lacey, and Sewell Peakman. A single autoimmune t cell receptor recognizes more than a million different peptides. Journal of Biological Chemistry, 287(92):1168–1177, 2011. doi: 10.1074/jbc.M111.289488. URL https://www.sciencedirect.com/science/article/pii/S0167569998012997.

[15] Amalie K Bentzen, Lina Such, Kamilla K Jensen, Andrea M Marquard, Leon E Jessen, Natalie J Miller, Candice D Church, Rikke Lyngaa, David M Koelle, Jürgen C Becker, Carsten Linnemann, Ton N M Schumacher, Paolo Marcatili, Paul Nghiem, Morten Nielsen, and Sine R Hadrup. T cell receptor fingerprinting enables in-depth characterization of the interactions governing recognition of peptide-mhc complexes. Nature biotechnology, November 2018. ISSN 1087-0156. doi: 10.1038/nbt.4303. URL https://europepmc.org/articles/PMC9452375.

[16] Andrew Sewell. Why must t cells be cross-reactive? Nature reviews. Immunology, 12:669–77, 08 2012. doi: 10.1038/nri3279.

[17] Dan Hudson, Ricardo A Fernandes, Mark Basham, Graham Ogg, and Hashem Koohy. Can we predict t cell specificity with digital biology and machine learning? Nature Reviews Immunology, pages 1–11, 2023.

[18] Jacob Glanville, Huang Huang, Allison Nau, Olivia Hatton, Lisa Wagar, Florian Rubelt, Xuhuai Ji, Arnold Han, Sheri Krams, Christina Pettus, Nikhil Haas, Cecilia Lindestam Arlehamn, Alessandro Sette, Scott Boyd, Thomas Scriba, Olivia Martinez, and Mark Davis. Identifying specificity groups in the t cell receptor repertoire. Nature, 547, 06 2017. doi: 10.1038/nature22976.

[19] Pradyot K Dash, Andrew J. Fiore-Gartland, Tomer Hertz, George C. Wang, Shalini Sharma, Aisha Souquette, Jeremy Chase Crawford, E Bridie Clemens, Thi H.O. Nguyen, Katherine Kedzierska, Nicole L. La Gruta, Philip Bradley, and Paul G. Thomas. Quantifiable predictive features define epitope-specific t cell receptor repertoires. Nature, 547(7661):89–93, July 2017. ISSN 0028-0836. doi: 10.1038/nature22383.

[20] Vanessa Isabell Jurtz, Leon Eyrich Jessen, Amalie Kai Bentzen, Martin Closter Jespersen, Swapnil Mahajan, Randi Vita, Kamilla Kjærgaard Jensen, Paolo Marcatili, Sine Reker Hadrup, Bjoern Peters, and Morten Nielsen. Nettcr: sequence-based prediction of tcr binding to peptide-mhc complexes using convolutional neural networks. bioRxiv, 2018. doi: 10.1101/433706. URL https://www.biorxiv.org/content/early/2018/10/02/433706.

[21] Pieter Moris, Joey De Pauw, Anna Postovskaya, Sofie Gielis, Nicolas De Neuter, Wout Bittremieux, Benson Ogunjimi, Kris Laukens, and Pieter Meysman. Current challenges for epitope-agnostic tcr interaction prediction and a new perspective derived from image classification. bioRxiv, 2020. doi: 10.1101/2019.12.18.880146. URL https://www.biorxiv.org/content/early/2020/09/11/2019.12.18.880146.

[22] Esteban Lanzarotti, Paolo Marcatili, and Morten Nielsen. Identification of the cognate peptide-mhc target of t cell receptors using molecular modeling and force field scoring. Molecular Immunology, 94:91–97, 2018. ISSN 0161-5890. doi: 10.1016/j.molimm.2017.12.019. URL https://www.sciencedirect.com/science/article/pii/S0161589017306211.

[23] Xingcheng Lin, Jason T. George, Nicholas P. Schafer, Kevin Ng Chau, Michael E. Birnbaum, Cecilia Clementi, José N. Onuchic, and Herbert Levine. Rapid assessment of t-cell receptor specificity of the immune repertoire. bioRxiv, 2021. doi: 10.1101/2020.04.06.028415. URL https://www.biorxiv.org/content/early/2021/04/22/2020.04.06.028415.

[24] Masato Ogishi and Hiroshi Yotsuyanagi. Quantitative prediction of the landscape of t cell epitope immunogenicity in sequence space. Frontiers in Immunology, 10, 2019. ISSN 1664-3224. doi: 10.3389/fimmu.2019.00827. URL https://www.frontiersin.org/articles/10.3389/fimmu.2019.00827.

[25] Martina Milighetti, John Shawe-Taylor, and Benny Chain. Predicting t cell receptor antigen specificity from structural features derived from homology models of receptor-peptide-major histocompatibility complexes. Frontiers in Physiology, 12, 2021. ISSN 1664-042X. doi: 10.3389/fphys.2021.730908. URL https://www.frontiersin.org/articles/10.3389/fphys.2021.730908.

[26] Yao Tong, Jiayin Wang, Tian Zheng, Xuanping Zhang, Xiao Xiao, Xiaoyan Zhu, Xin Lai, and Xiang Liu. Sete: Sequence-based ensemble learning approach for tcr epitope binding prediction. Computational Biology and Chemistry, 87:107281, 2020. ISSN 1476-9271. doi: 10.1016/j.compbiolchem.2020.107281. URL https://www.sciencedirect.com/science/article/pii/S1476927120303194.

[27] Ido Springer, Nili Tickotsky, and Yoram Louzoun. Contribution of t cell receptor alpha and beta cdr3, mhc typing, v and j genes to peptide binding prediction. Frontiers in Immunology, 12, 2021. ISSN 1664-3224. doi: 10.3389/fimmu.2021.664514. URL https://www.frontiersin.org/articles/10.3389/fimmu.2021.664514.

[28] Anna Weber, Jannis Born, and María Rodriguez Martínez. TITAN: T-cell receptor specificity prediction with bimodal attention networks. Bioinformatics, 37(Supplement):i237–i244, 07 2021. ISSN 1367-4803. doi: 10.1093/bioinformatics/btab294. URL 10.1093/bioinformatics/btab294.

[29] Yicheng Gao, Yuli Gao, Yuxiao Fan, Chengyu Zhu, Zhiting Wei, Chi Zhou, Guohui Chuai, Qinchang Chen, He Zhang, and Qi Liu. Pan-peptide meta learning for t-cell receptor–antigen binding recognition. Nature Machine Intelligence, 5:236–249, 2023.

[30] Bjørn P. Y. Kwee, Marius Messemaker, Eric Marcus, Giacomo Oliveira, Wouter Scheper, Catherine J. Wu, Jonas Teuwen, and Ton N. Schumacher. Stapler: Efficient learning of tcr-peptide specificity prediction from full-length tcr-peptide data. bioRxiv, 2023. doi: 10.1101/2023.04.25.538237. URL https://www.biorxiv.org/content/early/2023/04/28/2023.04.25.538237.

[31] Barthelemy Meynard-Piganeau, Christoph Feinauer, Martin Weigt, Aleksandra M. Walczak, and Thierry Mora. Tulip — a transformer based unsupervised language model for interacting peptides and t-cell receptors that generalizes to unseen epitopes. bioRxiv, 2023. doi: 10.1101/2023.07.19.549669. URL https://www.biorxiv.org/content/early/2023/07/19/2023.07.19.549669.

[32] Yi Han, Yuqiu Yang, Yanhua Tian, Farjana J. Fattah, Mitchell S. von Itzstein, Minying Zhang, Xiongbin Kang, Donghan M. Yang, Jialiang Liu, Yaming Xue, Chaoying Liang, Indu Raman, Chengsong Zhu, Olivia Xiao, Yifei Hu, Jonathan E. Dowell, Jade Homsi, Sawsan Rashdan, Shengjie Yang, Mary E. Gwin, David Hsiehchen, Yvonne Gloria-McCutchen, Ke Pan, Fangjiang Wu, Don Gibbons, Xinlei Wang, Cassian Yee, Junzhou Huang, Alexandre Reuben, Chao Cheng, Jianjun Zhang, David E. Gerber, and Tao Wang. pan-mhc and cross-species prediction of t cell receptor-antigen binding. bioRxiv, 2023. doi: 10.1101/2023.12.01.569599. URL https://www.biorxiv.org/content/early/2023/12/12/2023.12.01.569599.

[33] Yumeng Zhang, Zhikang Wang, Yunzhe Jiang, Dene R. Littler, Mark Gerstein, Anthony W. Purcell, Jamie Rossjohn, Hong-Yu Ou, and Jiangning Song. Epitope-anchored contrastive transfer learning for paired cd8+ t cell receptor–antigen recognition. Nature Machine Intelligence, Oct 2024. ISSN 2522-5839. doi: 10.1038/s42256-024-00913-8. URL 10.1038/s42256-024-00913-8.

[34] Kevin Wu, Kathryn E. Yost, Bence Daniel, Julia A. Belk, Yu Xia, Takeshi Egawa, Ansuman Satpathy, Howard Y. Chang, and James Zou. Tcr-bert: learning the grammar of t-cell receptors for flexible antigen-xbinding analyses. bioRxiv, 2021. doi: 10.1101/2021.11.18.469186. URL https://www.biorxiv.org/content/early/2021/11/20/2021.11.18.469186.

[35] Kristian Davidsen, Branden J Olson, III DeWitt, William S, Jean Feng, Elias Harkins, Philip Bradley, and IV Matsen, Frederick A. Deep generative models for t cell receptor protein sequences. eLife, 8:e46935, sep 2019. ISSN 2050-084X. doi: 10.7554/eLife.46935. URL 10.7554/eLife.46935.

[36] Giulio Isacchini, Aleksandra M. Walczak, Thierry Mora, and Armita Nourmohammad. Deep generative selection models of t and b cell receptor repertoires with sonnia. Proceedings of the National Academy of Sciences, 118(14):e2023141118, 2021. doi: 10.1073/pnas.2023141118. URL https://www.pnas.org/doi/abs/10.1073/pnas.2023141118.

[37] Ziqi Chen, Martin Renqiang Min, Hongyu Guo, Chao Cheng, Trevor Clancy, and Xia Ning. T-cell receptor optimization with reinforcement learning and mutation policies for precesion immunotherapy, 2023.

[38] Ethan Fast, Manjima Dhar, and Binbin Chen. Tapir: a t-cell receptor language model for predicting rare and novel targets. bioRxiv, 2023. doi: 10.1101/2023.09.12.557285. URL https://www.biorxiv.org/content/early/2023/09/15/2023.09.12.557285.

[39] Helder V. Ribeiro-Filho, Gabriel E. Jara, João V. S. Guerra, Melyssa Cheung, Nathaniel R. Felbinger, José G. C. Pereira, Brian G. Pierce, and Paulo S. Lopes de Oliveira. Exploring the potential of structure-based deep learning approaches for t cell receptor design. bioRxiv, 2024. doi: 10.1101/2024.04.19.590222. URL https://www.biorxiv.org/content/early/2024/04/24/2024.04.19.590222.

[40] Dhuvarakesh Karthikeyan, Colin Raffel, Benjamin Vincent, and Alex Rubinsteyn. Conditional generation of antigen specific t-cell receptor sequences. In NeurIPS 2023 Generative AI and Biology (GenBio) Workshop, 2023. URL https://openreview.net/forum?id=SckdgVW3Kq.

[41] Zhenghong Zhou, Junwei Chen, Shenggeng Lin, Liang Hong, Dong-Qing Wei, and Yi Xiong. Gratcr: epitope-specific t cell receptor sequence generation with data-efficient pre-trained models. bioRxiv, 2024. doi: 10.1101/2024.07.21.604503. URL https://www.biorxiv.org/content/early/2024/07/23/2024.07.21.604503.

[42] Yicheng Lin, Dandan Zhang, and Yun Liu. Tcr-gpt: Integrating autoregressive model and reinforcement learning for t-cell receptor repertoires generation, 2024. URL https://arxiv.org/abs/2408.01156.

[43] Mike Lewis, Yinhan Liu, Naman Goyal, Marjan Ghazvininejad, Abdelrahman Mohamed, Omer Levy, Ves Stoyanov, and Luke Zettlemoyer. Bart: Denoising sequence-to-sequence pre-training for natural language generation, translation, and comprehension, 2019.

[44] Colin Raffel, Noam Shazeer, Adam Roberts, Katherine Lee, Sharan Narang, Michael Matena, Yanqi Zhou, Wei Li, and Peter J. Liu. Exploring the limits of transfer learning with a unified text-to-text transformer, 2020.

[45] Barry Haddow, Rachel Bawden, Antonio Valerio Miceli Barone, Jindřich Helcl, and Alexandra Birch. Survey of Low-Resource Machine Translation. Computational Linguistics, 48(3):673–732, 09 2022. ISSN 0891-2017. doi: 10.1162/coli_a_00446. URL 10.1162/coli_a_00446.

[46] Rico Sennrich, Barry Haddow, and Alexandra Birch. Improving neural machine translation models with monolingual data, 2016.

[47] Junxian He, Jiatao Gu, Jiajun Shen, and Marc’Aurelio Ranzato. Revisiting self-training for neural sequence generation, 2020.

[48] Yinhan Liu, Jiatao Gu, Naman Goyal, Xian Li, Sergey Edunov, Marjan Ghazvininejad, Mike Lewis, and Luke Zettlemoyer. Multilingual denoising pre-training for neural machine translation, 2020.

[49] Jiajun Shen, Peng-Jen Chen, Matt Le, Junxian He, Jiatao Gu, Myle Ott, Michael Auli, and Marc’Aurelio Ranzato. The source-target domain mismatch problem in machine translation, 2020.

[50] Xing Niu, Michael Denkowski, and Marine Carpuat. Bi-directional neural machine translation with synthetic parallel data. In Alexandra Birch, Andrew Finch, Thang Luong, Graham Neubig, and Yusuke Oda, editors, Proceedings of the 2nd Workshop on Neural Machine Translation and Generation, pages 84–91, Melbourne, Australia, July 2018. Association for Computational Linguistics. doi: 10.18653/v1/W18-2710. URL https://aclanthology.org/W18-2710.

[51] Hsiu-Wei Yang, Yanyan Zou, Peng Shi, Wei Lu, Jimmy J. Lin, and Xu Sun. Aligning cross-lingual entities with multi-aspect information. In Conference on Empirical Methods in Natural Language Processing, 2019. URL https://api.semanticscholar.org/CorpusID:202121966.

[52] Liang Ding, Di Wu, and Dacheng Tao. Improving neural machine translation by bidirectional training. In Marie-Francine Moens, Xuanjing Huang, Lucia Specia, and Scott Wen-tau Yih, editors, Proceedings of the 2021 Conference on Empirical Methods in Natural Language Processing, pages 3278–3284, Online and Punta Cana, Dominican Republic, November 2021. Association for Computational Linguistics. doi: 10.18653/v1/2021.emnlp-main.263. URL https://aclanthology.org/2021.emnlp-main.263.

[53] Kun Yu, Ji Shi, Dan Lu, and Qiong Yang. Comparative analysis of CDR3 regions in paired human αβ CD8 T cells. FEBS Open Bio, 9(8):1450–1459, August 2019.

[54] Ilka Hoof, Bjoern Peters, John Sidney, Lasse Eggers Pedersen, Alessandro Sette, Ole Lund, Søren Buus, and Morten Nielsen. Netmhcpan, a method for mhc class i binding prediction beyond humans. Immunogenetics, 61:1–13, 2009.

[55] Edo Dotan, Gal Jaschek, Tal Pupko, and Yonatan Belinkov. Effect of tokenization on transformers for biological sequences. bioRxiv, 2023. doi: 10.1101/2023.08.15.553415. URL https://www.biorxiv.org/content/early/2023/08/17/2023.08.15.553415.

[56] Nili Tickotsky, Tal Sagiv, Jaime Prilusky, Eric Shifrut, and Nir Friedman. McPAS-TCR: a manually curated catalogue of pathology-associated T cell receptor sequences. Bioinformatics, 33(18):2924–2929, 05 2017. ISSN 1367-4803. doi: 10.1093/bioinformatics/btx286. URL 10.1093/bioinformatics/btx286.

[57] Mikhail Shugay, Dmitriy V Bagaev, Ivan V Zvyagin, Renske M Vroomans, Jeremy Chase Crawford, Garry Dolton, Ekaterina A Komech, Anastasiya L Sycheva, Anna E Koneva, Evgeniy S Egorov, Alexey V Eliseev, Ewald Van Dyk, Pradyot Dash, Meriem Attaf, Cristina Rius, Kristin Ladell, James E McLaren, Katherine K Matthews, E Bridie Clemens, Daniel C Douek, Fabio Luciani, Debbie van Baarle, Katherine Kedzierska, Can Kesmir, Paul G Thomas, David A Price, Andrew K Sewell, and Dmitriy M Chudakov. VDJdb: a curated database of T-cell receptor sequences with known antigen specificity. Nucleic Acids Research, 46(D1):D419–D427, 09 2017. ISSN 0305-1048. doi: 10.1093/nar/gkx760. URL 10.1093/nar/gkx760.

[58] Randi Vita, Swapnil Mahajan, James A Overton, Sandeep Kumar Dhanda, Sheridan Martini, Jason R Cantrell, Daniel K Wheeler, Alessandro Sette, and Bjoern Peters. The Immune Epitope Database (IEDB): 2018 update. Nucleic Acids Research, 47(D1):D339–D343, 10 2018. ISSN 0305-1048. doi: 10.1093/nar/gky1006. URL 10.1093/nar/gky1006.

[59] Jennifer N. Dines, Thomas J. Manley, Emily Svejnoha, Heidi M. Simmons, Ruth Taniguchi, Mark Klinger, Lance Baldo, and Harlan Robins. The immunerace study: A prospective multicohort study of immune response action to covid-19 events with the immunecode™ open access database. medRxiv, 2020. doi: 10.1101/2020.08.17.20175158. URL https://www.medrxiv.org/content/early/2020/08/21/2020.08.17.20175158.1.

[60] Timothy J. O’Donnell, Alex Rubinsteyn, and Uri Laserson. Mhcflurry 2.0: Improved pan-allele prediction of mhc class i-presented peptides by incorporating antigen processing. Cell Systems, 11(1):42–48.e7, 2020. ISSN 2405-4712. doi: 10.1016/j.cels.2020.06.010. URL https://www.sciencedirect.com/science/article/pii/S2405471220302398.

[61] Katherine Lee, Daphne Ippolito, Andrew Nystrom, Chiyuan Zhang, Douglas Eck, Chris Callison-Burch, and Nicholas Carlini. Deduplicating training data makes language models better, 2022. URL https://arxiv.org/abs/2107.06499.

[62] Asa Cooper Stickland, Xian Li, and Marjan Ghazvininejad. Recipes for adapting pre-trained monolingual and multilingual models to machine translation. In Proceedings of the 16th Conference of the European Chapter of the Association for Computational Linguistics: Main Volume. Association for Computational Linguistics, 2021. doi: 10.18653/v1/2021.eacl-main.301. URL 10.18653/v1/2021.eacl-main.301.

[63] Barret Zoph, Deniz Yuret, Jonathan May, and Kevin Knight. Transfer learning for low-resource neural machine translation. ArXiv, abs/1604.02201, 2016. URL https://api.semanticscholar.org/CorpusID:16631020.

[64] Si-Yi Chen, Tao Yue, Qian Lei, and An-Yuan Guo. TCRdb: a comprehensive database for T-cell receptor sequences with powerful search function. Nucleic Acids Research, 49 (D1):D468–D474, 09 2020. ISSN 0305-1048. doi: 10.1093/nar/gkaa796. URL 10.1093/nar/gkaa796.

[65] Ahmed Elnaggar, Michael Heinzinger, Christian Dallago, Ghalia Rihawi, Yu Wang, Llion Jones, Tom Gibbs, Tamas Feher, Christoph Angerer, Debsindhu Bhowmik, and Burkhard Rost. Prottrans: Towards cracking the language of life’s code through self-supervised deep learning and high performance computing. bioRxiv, 2020. doi: 10.1101/2020.07.12.199554. URL https://www.biorxiv.org/content/early/2020/07/12/2020.07.12.199554.

[66] Kaitao Song, Xu Tan, Tao Qin, Jianfeng Lu, and Tie-Yan Liu. Mass: Masked sequence to sequence pre-training for language generation, 2019.

[67] Guillaume Lample and Alexis Conneau. Cross-lingual language model pretraining, 2019.

[68] Ilya Sutskever, Oriol Vinyals, and Quoc V. Le. Sequence to sequence learning with neural networks, 2014.

[69] Kyunghyun Cho, Bart van Merrienboer, Caglar Gulcehre, Dzmitry Bahdanau, Fethi Bougares, Holger Schwenk, and Yoshua Bengio. Learning phrase representations using rnn encoder-decoder for statistical machine translation, 2014.

[70] Bryan Eikema and Wilker Aziz. Is map decoding all you need? the inadequacy of the mode in neural machine translation, 2020.

[71] Thibault Sellam, Dipanjan Das, and Ankur P. Parikh. Bleurt: Learning robust metrics for text generation, 2020. URL https://arxiv.org/abs/2004.04696.

[72] Ricardo Rei, Craig Stewart, Ana C Farinha, and Alon Lavie. Comet: A neural framework for mt evaluation, 2020. URL https://arxiv.org/abs/2009.09025.

[73] Filippo Grazioli, Anja Mösch, Pierre Machart, Kai Li, Israa Alqassem, Timothy J O’Donnell, and Martin Renqiang Min. On tcr binding predictors failing to generalize to unseen peptides. Frontiers in Immunology, 13:1014256, 2022.

[74] Morten Nielsen, Anne Eugster, Mathias Fynbo Jensen, Manisha Goel, Andreas Tiffeau-Mayer, Aurelien Pelissier, Sebastiaan Valkiers, María Rodríguez Martínez, Barthélémy Meynard-Piganeeau, Victor Greiff, Thierry Mora, Aleksandra M. Walczak, Giancarlo Croce, Dana L. Moreno, David Gfeller, Pieter Meysman, and Justin Barton. Lessons learned from the immrep23 tcr-epitope prediction challenge. ImmunoInformatics, 16, Dec 2024. ISSN 2667-1190. doi: 10.1016/j.immuno.2024.100045. URL 10.1016/j.immuno.2024.100045.

[75] Herman N Eisen and Gregory W Siskind. Variations in affinities of antibodies during the immune response. Biochemistry, 3(7):996–1008, 1964.

[76] Sharrol Bachas, Goran Rakocevic, David Spencer, Anand V. Sastry, Robel Haile, John M. Sutton, George Kasun, Andrew Stachyra, Jahir M. Gutierrez, Edriss Yassine, Borka Medjo, Vincent Blay, Christa Kohnert, Jennifer T. Stanton, Alexander Brown, Nebojsa Tijanic, Cailen McCloskey, Rebecca Viazzo, Rebecca Consbruck, Hayley Carter, Simon Levine, Shaheed Abdulhaqq, Jacob Shaul, Abigail B. Ventura, Randal S. Olson, Engin Yapici, Joshua Meier, Sean McClain, Matthew Weinstock, Gregory Hannum, Ariel Schwartz, Miles Gander, and Roberto Spreafico. Antibody optimization enabled by artificial intelligence predictions of binding affinity and naturalness. bioRxiv, 2022. doi: 10.1101/2022.08.16.504181. URL https://www.biorxiv.org/content/early/2022/08/17/2022.08.16.504181.

[77] Kevin Michalewicz, Mauricio Barahona, and Barbara Bravi. Antipasti: interpretable prediction of antibody binding affinity exploiting normal modes and deep learning. bioRxiv, 2023. doi: 10.1101/2023.12.22.572853. URL https://www.biorxiv.org/content/early/2023/12/23/2023.12.22.572853.

[78] Nishant Kumar Singh, Timothy P. Riley, Sarah Catherine B. Baker, Tyler Borrman, Zhiping Weng, and Brian M. Baker. Emerging concepts in tcr specificity: Rationalizing and (maybe) predicting outcomes. The Journal of Immunology, 199:2203–2213, 2017. URL https://api.semanticscholar.org/CorpusID:5375575.

[79] K. Christopher Garcia, Massimo Degano, Larry R. Pease, Mingdong Huang, Per A. Peterson, Luc Teyton, and Ian A. Wilson. Structural basis of plasticity in t cell receptor recognition of a self peptide-mhc antigen. Science, 279(5354):1166–1172, 1998. doi: 10.1126/science.279.5354.1166. URL https://www.science.org/doi/abs/10.1126/science.279.5354.1166.

[80] Xiang Zhao, Elizabeth M. Kolawole, Waipan Chan, Yinnian Feng, Xinbo Yang, Marvin H. Gee, Kevin M. Jude, Leah V. Sibener, Polly M. Fordyce, Ronald N. Germain, Brian D. Evavold, and K. Christopher Garcia. Tuning t cell receptor sensitivity through catch bond engineering. Science, 376(6589):eabl5282, 2022. doi: 10.1126/science.abl5282. URL https://www.science.org/doi/abs/10.1126/science.abl5282.

[81] Kishore Papineni, Salim Roukos, Todd Ward, and Wei-Jing Zhu. Bleu: A method for automatic evaluation of machine translation. In Proceedings of the 40th Annual Meeting on Association for Computational Linguistics, ACL ‘02, page 311–318, USA, 2002. Association for Computational Linguistics. doi: 10.3115/1073083.1073135. URL 10.3115/1073083.1073135.

[82] Zachary Sethna, Yuval Elhanati, Jr Callan, Curtis G, Aleksandra M Walczak, and Thierry Mora. OLGA: fast computation of generation probabilities of B- and T-cell receptor amino acid sequences and motifs. Bioinformatics, 35(17):2974–2981, 01 2019. ISSN 1367-4803. doi: 10.1093/bioinformatics/btz035. URL 10.1093/bioinformatics/btz035.

[83] Anirban Sarkar, Ziqi Tang, Chris Zhao, and Peter K Koo. Designing dna with tunable regulatory activity using discrete diffusion. bioRxiv, 2024. doi: 10.1101/2024.05.23.595630. URL https://www.biorxiv.org/content/early/2024/05/24/2024.05.23.595630.

[84] Kai W Wucherpfennig, Paul M Allen, Franco Celada, Irun R Cohen, Rob De Boer, K Christopher Garcia, Byron Goldstein, Ralph Greenspan, David Hafler, Philip Hodgkin, Erik S Huseby, David C Krakauer, David Nemazee, Alan S Perelson, Clemencia Pinilla, Roland K Strong, and Eli E Sercarz. Polyspecificity of T cell and B cell receptor recognition. Semin. Immunol., 19(4): 216–224, August 2007.

[85] Melvin Cohn. An in depth analysis of the concept of “polyspecificity” assumed to characterize TCR/BCR recognition. Immunol. Res., 40(2):128–147, 2008.

[86] Valentin Quiniou, Pierre Barennes, Vanessa Mhanna, Paul Stys, Helene Vantomme, Zhicheng Zhou, Federica Martina, Nicolas Coatnoan, Michele Barbie, Hang-Phuong Pham, Béatrice Clémenceau, Henri Vie, Mikhail Shugay, Adrien Six, Barbara Brandao, Roberto Mallone, Encarnita Mariotti-Ferrandiz, and David Klatzmann. Human thymopoiesis produces polyspecific CD8+ α/β T cells responding to multiple viral antigens. Elife, 12, March 2023.

[87] James Henderson, Yuta Nagano, Martina Milighetti, and Andreas Tiffeau-Mayer. Limits on inferring t-cell specificity from partial information, 2024. URL https://arxiv.org/abs/2404.12565.

[88] Pranav Rajpurkar, Jian Zhang, Konstantin Lopyrev, and Percy Liang. Squad: 100,000+ questions for machine comprehension of text, 2016.

[89] Alex Wang, Amanpreet Singh, Julian Michael, Felix Hill, Omer Levy, and Samuel R. Bowman. Glue: A multi-task benchmark and analysis platform for natural language understanding, 2019.

[90] Jacob Devlin, Ming-Wei Chang, Kenton Lee, and Kristina Toutanova. Bert: Pre-training of deep bidirectional transformers for language understanding, 2019.

[91] Alec Radford and Karthik Narasimhan. Improving language understanding by generative pre-training. 2018. URL https://api.semanticscholar.org/CorpusID:49313245.

[92] Zihao Fu, Wai Lam, Qian Yu, Anthony Man-Cho So, Shengding Hu, Zhiyuan Liu, and Nigel Collier. Decoder-only or encoder-decoder? interpreting language model as a regularized encoder-decoder, 2023.

[93] Ashish Vaswani, Noam Shazeer, Niki Parmar, Jakob Uszkoreit, Llion Jones, Aidan N. Gomez, Lukasz Kaiser, and Illia Polosukhin. Attention is all you need, 2023.

[94] Andreas Mayer and Curtis G. Callan. Measures of epitope binding degeneracy from t cell receptor repertoires. Proceedings of the National Academy of Sciences, 120(4):e2213264120, 2023. doi: 10.1073/pnas.2213264120. URL https://www.pnas.org/doi/abs/10.1073/pnas.2213264120.

[95] Don Mason. A very high level of crossreactivity is an essential feature of the t-cell receptor. Immunology Today, 19(9):395–404, 1998. ISSN 0167-5699. doi: 10.1016/S0167-5699(98)01299-7. URL https://www.sciencedirect.com/science/article/pii/S0167569998012997.

[96] Yuta Nagano and Benjamin Chain. tidytcells: standardizer for tr/mh nomenclature. Frontiers in Immunology, 14, 2023. ISSN 1664-3224. doi: 10.3389/fimmu.2023.1276106. URL https://www.frontiersin.org/journals/immunology/articles/10.3389/fimmu.2023.1276106.

[97] Niklas Muennighoff, Alexander Rush, Boaz Barak, Teven Le Scao, Nouamane Tazi, Aleksandra Piktus, Sampo Pyysalo, Thomas Wolf, and Colin A Raffel. Scaling data-constrained language models. In A. Oh, T. Naumann, A. Globerson, K. Saenko, M. Hardt, and S. Levine, editors, Advances in Neural Information Processing Systems, volume 36, pages 50358–50376. Curran Associates, Inc., 2023. URL https://proceedings.neurips.cc/paper_files/paper/2023/file/9d89448b63ce1e2e8dc7af72c984c196-Paper-Conference.pdf.

[98] Dumitru Erhan, Yoshua Bengio, Aaron Courville, Pierre-Antoine Manzagol, Pascal Vincent, and Samy Bengio. Why does unsupervised pre-training help deep learning? J. Mach. Learn. Res., 11:625–660, March 2010. ISSN 1532-4435.

